# Molecular pathways and cellular subsets associated with adverse clinical outcomes in overlapping immune-related myocarditis and myositis

**DOI:** 10.1101/2023.09.15.556590

**Authors:** Bilal A. Siddiqui, Nicolas L. Palaskas, Sreyashi Basu, Yibo Dai, Zhong He, Shalini S. Yadav, James P. Allison, Rahul Sheth, Sudhakar Tummala, L. Maximilian Buja, Meenakshi Bhattacharjee, Cezar A. Iliescu, Anishia Rawther-Karedath, Anita Deswal, Linghua Wang, Padmanee Sharma, Sumit K. Subudhi

## Abstract

Immune checkpoint therapies (ICTs) can induce life-threatening immune-related adverse events, including myocarditis and myositis, which are rare but often concurrent. The molecular pathways and immune subsets underlying these toxicities remain poorly understood. To address this need, we obtained heart and skeletal muscle biopsies for single-cell RNA sequencing in living patients with cancers treated with ICTs admitted to the hospital with myocarditis and /or myositis (overlapping myocarditis plus myositis, n=10; myocarditis-only, n=1) compared to ICT-exposed patients ruled out for toxicity utilized as controls (n=9) within 96 hours of clinical presentation. Analyses of 58,523 cells revealed clonally expanded CD8^+^ T cells with a cytotoxic phenotype expressing activation/exhaustion markers in both myocarditis and myositis. Furthermore, the analyses identified a population of tissue-resident myeloid cells expressed Fc_γ_RIIIa, which is known to bind IgG and regulate complement activation. Immunohistochemistry of affected cardiac and skeletal muscle tissues revealed protein expression of pan-IgG and complement product C4d that were associated with the presence of high-titer serum autoantibodies against muscle antigens in a subset of patients. We further identified a population of inflammatory IL-1B^+^TNF^+^ myeloid cells specifically enriched in myocarditis and associated with greater toxicity severity and poorer clinical outcomes. These results are the first to recognize these myeloid subsets in human immune-related myocarditis and myositis tissues and nominate new targets for investigation into rational treatments to overcome these high-mortality toxicities.

**Graphical Abstract:** 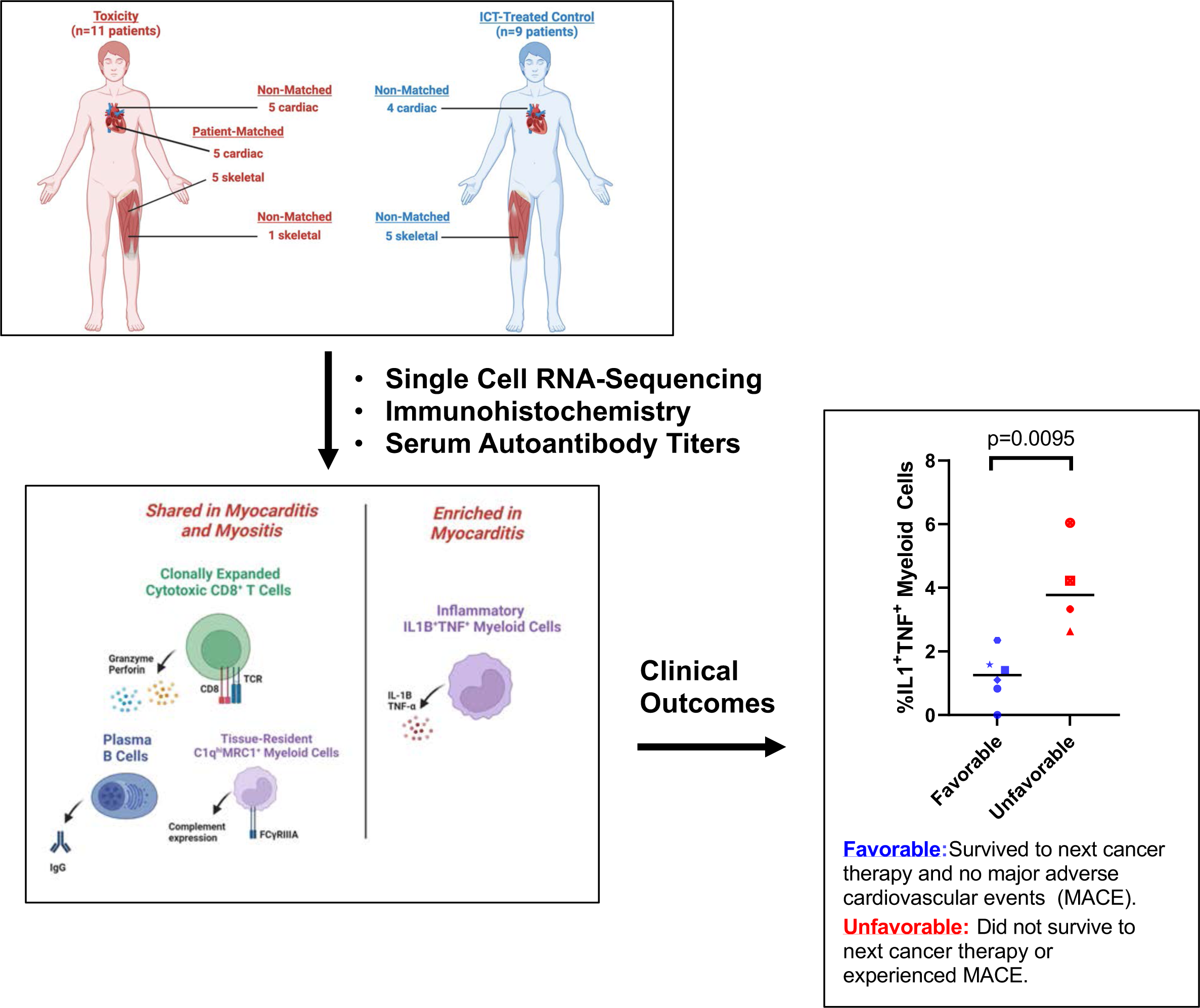

## Introduction

Immune checkpoint therapies (ICTs) induce anti-tumor T cell responses and long-term survival in a subset of patients. However, ICTs can also trigger life-threatening autoimmunity, termed immune-related adverse events (irAEs), that often require ICT discontinuation. Immune-related myocarditis, inflammation of the heart, is a rare irAE affecting approximately 1% of patients and carries up to 40% mortality with standard first-line treatment with corticosteroids.^1^ This often co-occurs with immune-related myositis (inflammation of skeletal muscle) and myasthenia gravis (MG, antibody-mediated disruption of neuromuscular transmission). The molecular pathways and immune subsets underlying these irAEs are incompletely understood and remain a critical unmet need for developing rational immunosuppressive treatment strategies to ameliorate these toxicities.

The first report of fatal immune-related myocarditis and myositis reported intracardiac infiltration of CD4^+^ and CD8^+^ T cells and CD68^+^ macrophages in patient autopsy specimens, with shared T cell receptor (TCR) clones among heart, skeletal muscle, and tumor, suggesting that antigen cross-reactivity may contribute to pathogenesis.^2^ Consistent with these findings, α-myosin-expanded TCRs were identified in cardiac and skeletal muscle of patients with myocarditis, further supporting the role of antigen-specific CD8^+^ T cells in disease pathogenesis.^3^

Nonetheless, T cells do not function in isolation, requiring interaction of TCRs with MHC-bound peptide antigens expressed on antigen-presenting cells (APCs) and costimulatory signals for optimal activation, differentiation, and proliferation. Myeloid cells, which encompass diverse cell types including macrophages and dendritic cells (DCs) that can fulfill these functions, have been shown to contribute to irAEs such as colitis and arthritis.^4,5^ However, their role in myocarditis and myositis remains undefined, despite their significant presence in inflamed muscle tissues.^4–6^ Herein, we characterized the cardiac and skeletal muscle tissue microenvironments utilizing single-cell transcriptomics from biopsy samples obtained from patients hospitalized with clinically-proven, overlapping immune-related myocarditis and myositis. In parallel, we performed comparative analyses on similar tissues from patients who were clinically ruled out for immune-related myocarditis and myositis to control for the effects of ICT exposure. These studies were complemented by histopathologic analyses and evaluation of serum immunoglobulins. Optimization of biopsy techniques allowed the safe and rapid collection of fresh heart and skeletal muscle tissues in living, severely ill patients early in their disease course. Overall, we identified candidate molecular pathways and immune subpopulations in immune-related myocarditis and myositis for investigation into rational immunosuppressive therapeutic strategies.

## Results

### Patient characteristics and tissue specimens

We collected clinical data and tissue samples from 20 living patients with cancers treated with ICTs and admitted to the hospital with suspected immune-related myocarditis and/or myositis, who had evaluable tissue for further analysis **(Table 1) (Supplementary Tables 1 & 2)**. A total of 11 patients were confirmed to have an irAE diagnosis (myocarditis plus myositis, n=10; myocarditis-only, n=1 [patient 4388]) **(Table 1) (Supplementary Table 1) (Supplementary Figure 1a)**. The 9 patients that were treated with ICT and clinically ruled out for immune-related myocarditis and myositis were utilized as controls (cardiac muscle, n=4; skeletal muscle, n=5) **(Table 1) (Supplementary Table 2)**.^7,8^ Serum markers of tissue injury were consistent with the adjudicated diagnoses. The median serum levels of troponin-T, a component of the troponin complex of proteins integral to contraction of skeletal and cardiac muscles and commonly utilized as a measurement of cardiac tissue damage, were 1,056 ng/mL in the confirmed irAE group (95% CI: 485-4,613) and 60 ng/mL in the control group (95% CI: 19-358) (reference range: <18 ng/mL, p<0.0001) **(Supplementary Figure 1b)**. Similarly, the median levels of creatine kinase (CK), a protein that catalyzes conversion of phosphocreatine that is used as an energy reservoir in tissues that rapidly consume adenosine triphosphate (ATP) such as skeletal muscle and is a measure of muscle damage, were 1,373 U/L in the confirmed irAE group (95% CI: 182-6,012) and 235 U/L in the control group (95% CI: 59-717) (reference range: <180 U/L, p=0.0097) **(Supplementary Figure 1c).** The most common cancers in this patient cohort were renal cell carcinoma (RCC) and melanoma **(Table 1).** Although all patients received anti-PD-(L)1-based therapy, the most common ICT received in patients with confirmed irAEs was the combination of anti-CTLA-4 plus anti-PD-(L)1 (n=4, 36%) **(Table 1).** The median number of cycles of ICT in the confirmed irAEs group was 1 (interquartile range [IQR]: 1-2) and in the control group was 4 (IQR: 3-8) **(Table 1)**.

**Table 1:**
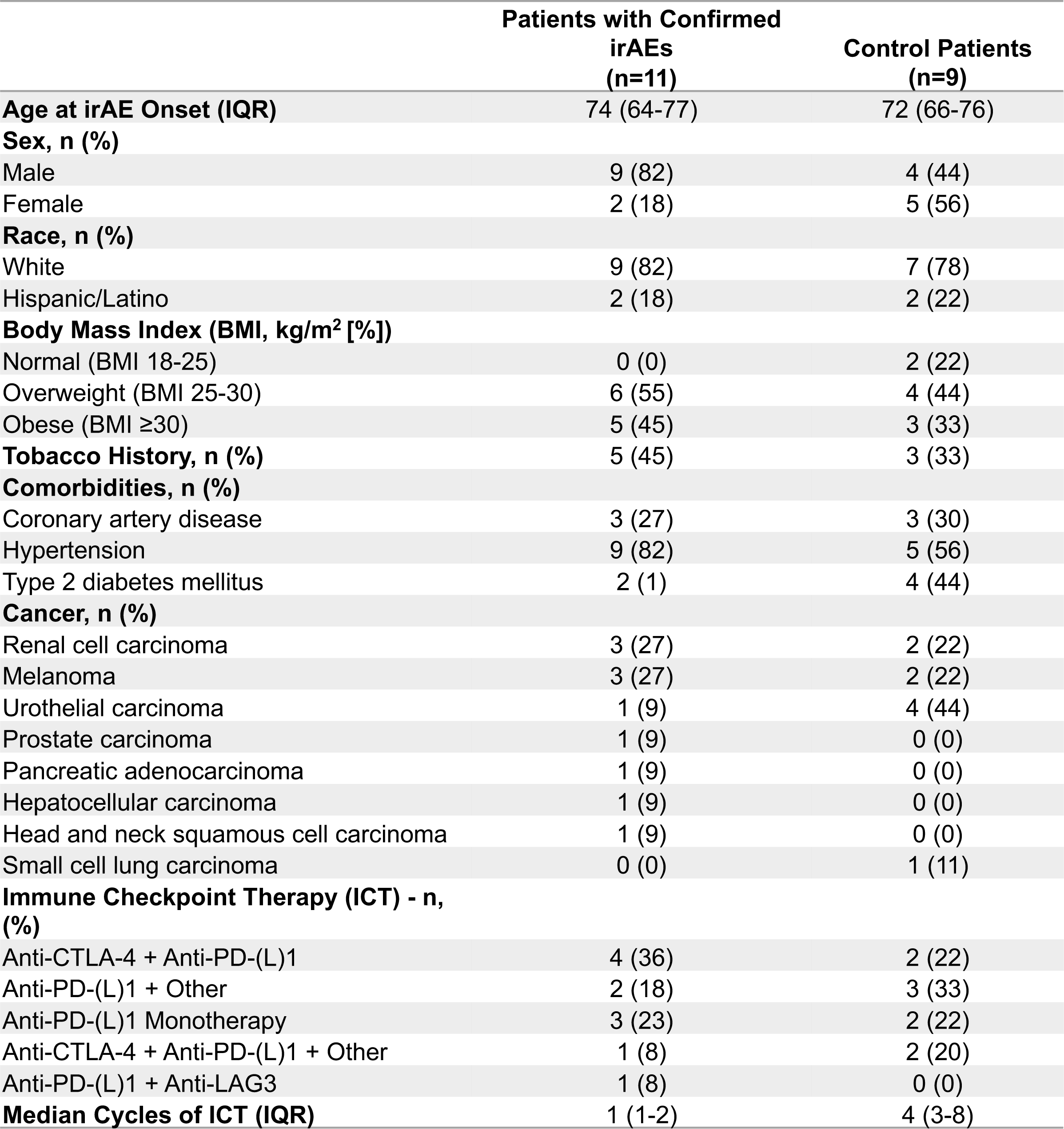
Baseline Clinical Characteristics.

### Overall cellular landscape of immune-related myocarditis and myositis

Given that immune-related myocarditis and myositis are rare, often concurrently occurring irAEs both involving muscle tissues, we surmised that they may also share molecular pathways and immune cell subsets in contributing to disease pathogenesis. Thus, to comprehensively define the phenotype and putative function of immune cells within the inflamed tissue microenvironment, we performed single-cell RNA sequencing (scRNA-seq) on cardiac (n=10 myocarditis; n=4 control) and skeletal muscle tissues (n=6 myositis; n=5 control) **(Figure 1a)**. Of the 11 total patients with confirmed irAEs, 10 patients had overlapping myocarditis and myositis, and 1 patient (4388) had confirmed myocarditis only (**Supplementary Table 1** & **Supplementary Figure 1a**). Paired patient-matched cardiac and skeletal muscle tissues were obtained in five patients **(Figure 1a).** A total of 27,594 live cells were evaluable for further analyses, with 4,591 live cells from normal cardiac tissue and 23,003 live cells from myocarditis tissue **(Figures 1b and 1c, respectively)**. From skeletal muscle (quadriceps), a total of 30,929 live cells were obtained, with 14,561 live cells from normal skeletal muscle and 16,368 live cells from myositis tissue **(Figures 1d and 1e, respectively)**. To ensure the evaluation of cell subsets was not skewed by differing cell numbers in toxicity versus control tissues, we downsampled the cells in the toxicity groups to match the number of cells in each control group, and visualized them under uniform manifold approximation and projection (UMAP) embeddings **(Supplementary Figure 2a).**

**Figure 1:**
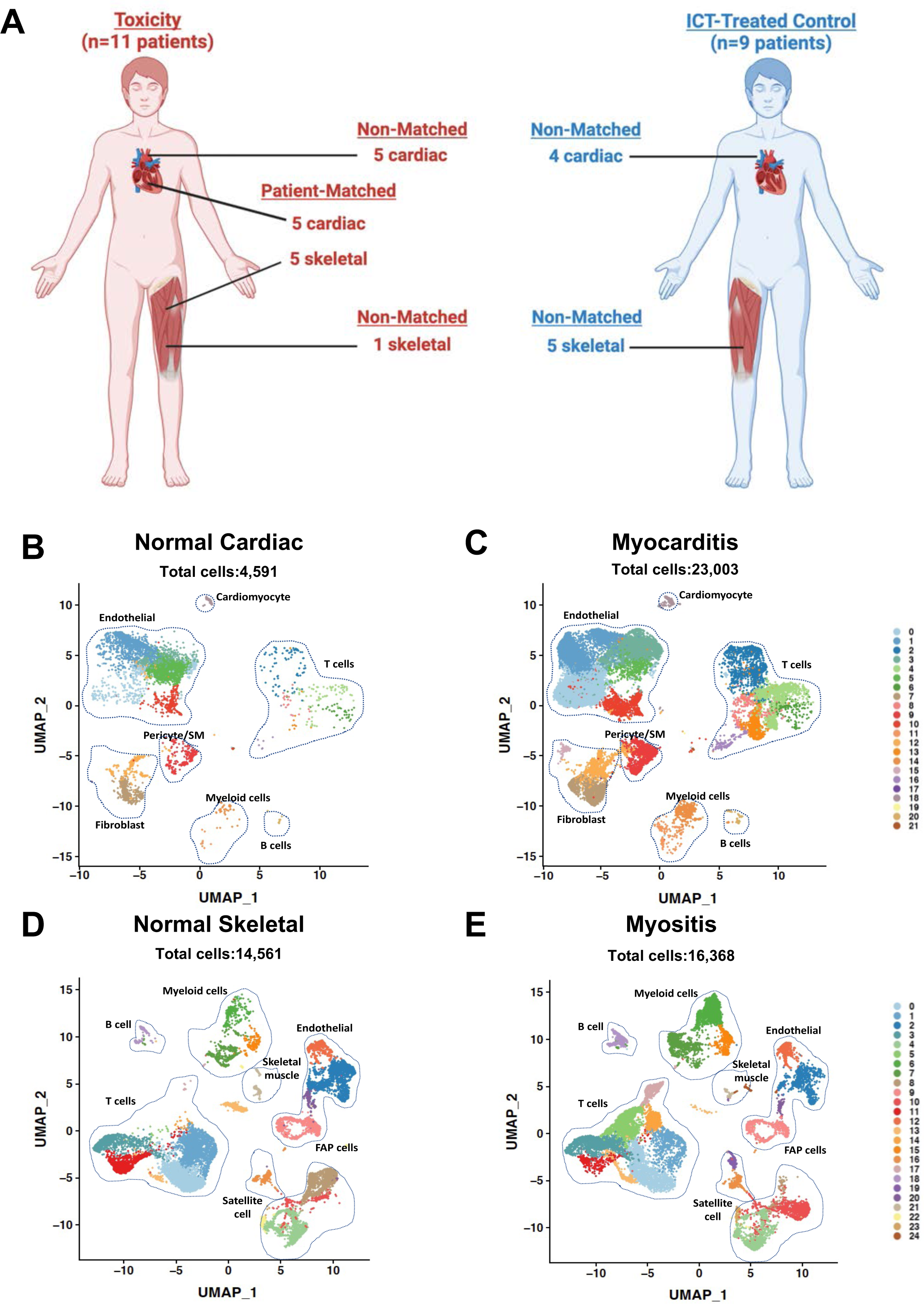
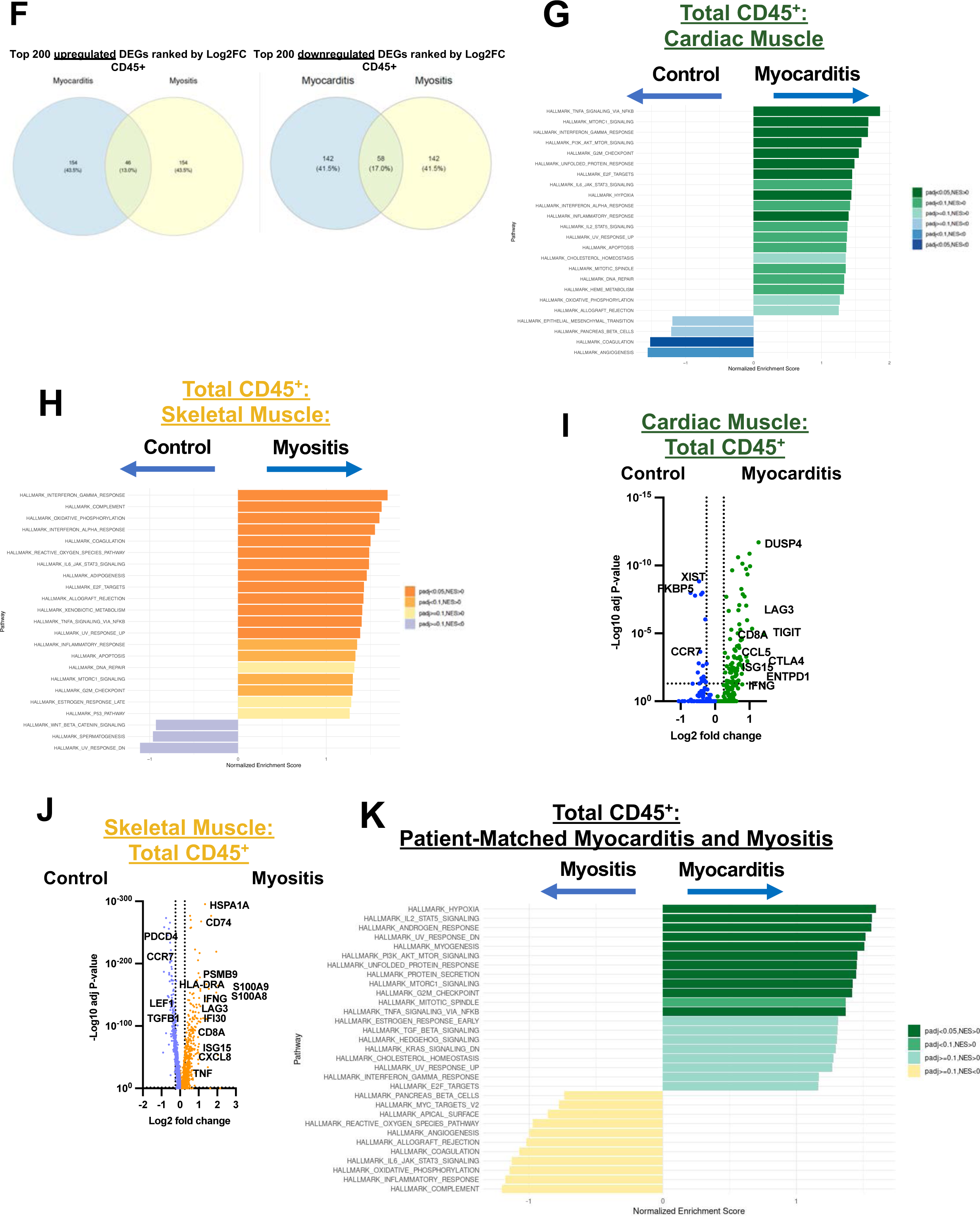
Overall cellular landscape of immune checkpoint therapy (ICT)-induced myocarditis and myositis. **A.** Overview of enrolled patients and collected specimens. **B.** Uniform Manifold Approximation and Projection (UMAP) plot of live cells obtained from normal cardiac muscle tissue. **C.** UMAP plot of live cells from myocarditis tissue. **D.** UMAP plot of live cells from normal skeletal muscle tissue. **E.** UMAP plot of live cells myositis tissue. **F.** Top upregulated and downregulated differentially expressed genes in CD45^+^ cells in myocarditis versus myositis samples. **G.** Differentially expressed hallmark transcriptional pathways in total CD45^+^ immune cells between myocarditis and control cardiac muscle tissue. **H.** Differentially expressed hallmark transcriptional pathways in total CD45^+^ immune cells between myositis and control skeletal muscle tissue. **I.** Volcano plot of differentially expressed genes in CD45+ immune cells in cardiac muscle. **J.** Volcano plot of differentially expressed genes in CD45+ immune cells in skeletal muscle. **K.** Differentially expressed hallmark transcriptional pathways in patient-matched myocarditis and myositis tissue.

To ensure that the biopsy methods obtained representative cardiac and skeletal muscle samples in both the toxicity and control groups, we first evaluated the CD45^-^ non-immune cell compartment. Within cardiac muscle, we obtained 22,411 cells consisting of cardiomyocytes, endothelial cells, fibroblasts, and pericytes/smooth muscle **(Supplementary Figures 2b & 2c).** Within skeletal muscle, we obtained 12,762 CD45^-^ cells that were comprised of skeletal muscle cells, endothelial cells, satellite cells (muscle stem cells), fibro-adipogenic precursors (FAPs), and pericytes/smooth muscle, which have previously been described in single-cell atlases of human skeletal muscle **(Supplementary Figures 2d & 2e).**^9^ Within these subpopulations, there were no difference in frequencies between control and immune-related myocarditis tissues **(Supplementary Figure 3)** nor between control and immune-related myositis samples **(Supplementary Figure 4)**. To formally compare the CD45^-^ cell types in the cardiac and skeletal muscle samples, we performed hierarchical clustering based on Euclidean distance of expression, which demonstrated as expected greater similarities for smooth muscle and endothelial cell clusters than for fibroblast and FAP subsets between cardiac and skeletal muscles **(Supplementary Figure 5).**

We next turned our attention to the CD45^+^ immune cell compartment (n=5,183 CD45^+^ cells in cardiac muscle samples; n=18,167 CD45^+^ cells in skeletal muscle samples) **(Figures 1b-1e)**. There was a trend towards a higher frequency of CD45^+^ cells in both immune-related myocarditis and myositis specimens compared to control samples **(Supplementary Figure 6)**. It should be noted that most patients received at least one dose of intravenous corticosteroids prior to biopsy, which may have attenuated the frequency of live immune cell populations within the inflamed muscles **(Table 1).** To evaluate the transcriptional profiles of CD45^+^ immune cells in inflamed muscle tissues, we quantified the top upregulated and downregulated differentially expressed genes between myocarditis and myositis samples and observed that a minority of genes were shared **(Figure 1f).** To determine the identity of these transcriptional differences, we performed gene set enrichment analysis (GSEA) on all pooled CD45^+^ immune cells. Within both cardiac and skeletal muscle, the TNF-α signalling via NF-κB and IFN-γ response pathways were observed to be highly upregulated, in addition to the complement pathway in skeletal muscle **(Figures 1g-h).** Consistent with these findings, genes associated with cytotoxic T cells (*CD8A*), T cell activation (*LAG3*), type I interferon (*ISG15*), and type II interferon (*IFNG*) were significantly differentially expressed in CD45^+^ cells in ICT-induced myocarditis and myositis compared to controls **(Figures 1i-j)**. To account for individual patient variation and isolate transcriptional pathways within CD45^+^ immune cells that were differentially expressed in myocarditis versus myositis, we directly compared patient-matched myocarditis and myositis samples **(Figure 1k)**. The top pathways differentially expressed in myocarditis meeting a padj<0.05 threshold included hallmark hypoxia and IL2/STAT5 signaling pathways; **(Figure 1k).**

### Immune cell subsets in ICT-induced myocarditis and myositis

To evaluate the overall immune cell populations in myocarditis and myositis, we compared the frequency of total lymphoid (T cell, NK cell, or B cell) and myeloid cell frequency within CD45^+^ cells in myocarditis versus control cardiac muscle samples and myositis and control skeletal muscle samples **(Supplementary Figures 6a and 6b, respectively),** and there were no observed differences in frequency. Therefore, we subclassified individual immune cell clusters and identified distinct clusters of CD8^+^ T cells (cytotoxic, *GZMK*-expressing, and proliferating), CD4^+^ T cells (naïve/central memory, T_reg_, cytotoxic), NK cells, naïve B cells, and myeloid cells (including SELL^+^ [L-selectin] macrophages, IL1B^+^ TNF^+^ myeloid cells, and conventional type 2 dendritic cells [cDC2s], professional APCs that prime CD4^+^ T cells) in cardiac muscle **(Figures 2a-2c).** Similarly, within skeletal muscle, we identified clusters of CD8^+^ T cells (cytotoxic, proliferating, and *GZMK*-expressing), CD4^+^ T cells (naïve/central memory), NK cells, and myeloid cells (including SELL^+^ myeloid, IL1B^+^TNF^+^ myeloid, and cDC2 populations **(Figures 2d-f)**. To formally compare the CD45^+^ cell types in the cardiac and skeletal muscle samples, we performed hierarchical clustering based on Euclidean distance of expression and observed shared clustering of the majority of specific cell subtypes in cardiac and skeletal muscle **(Figure 2g & Supplementary Figure 6).** Consistent with this observation, pooling of patient-matched myocarditis and myositis samples revealed overall similarity of the immune cell subclusters in both compartments **(Figure 2h-i).**

**Figure 2:**
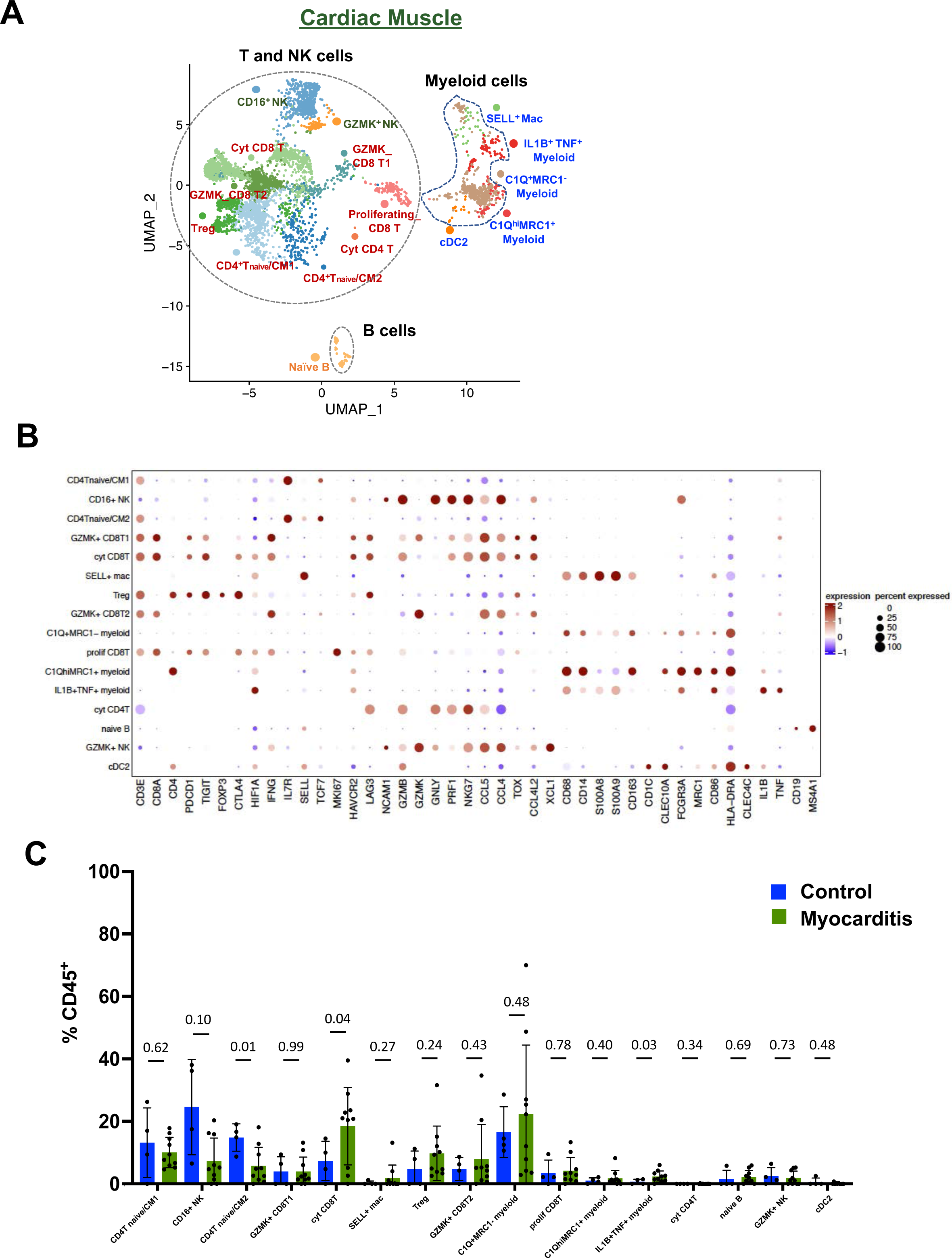
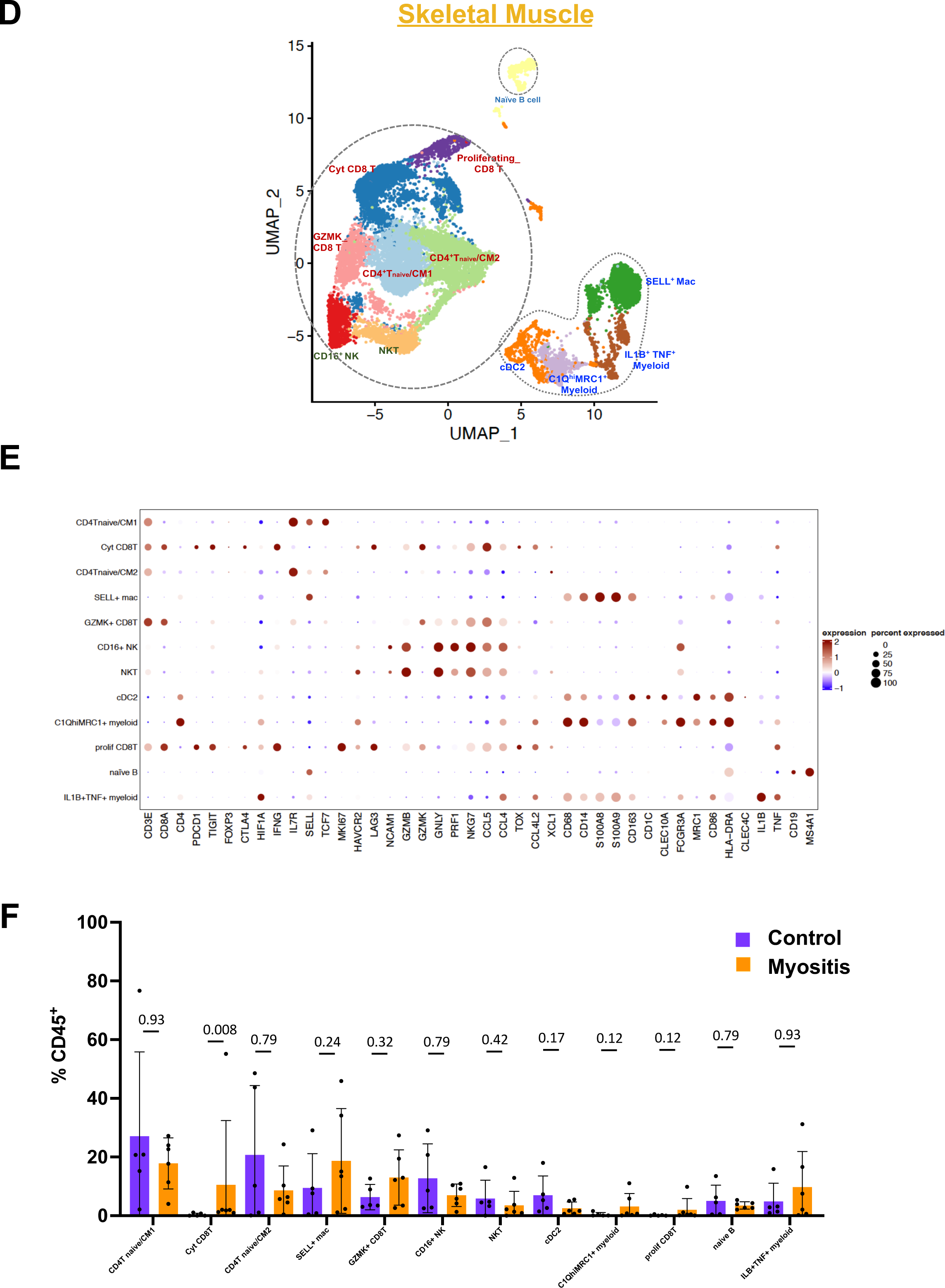
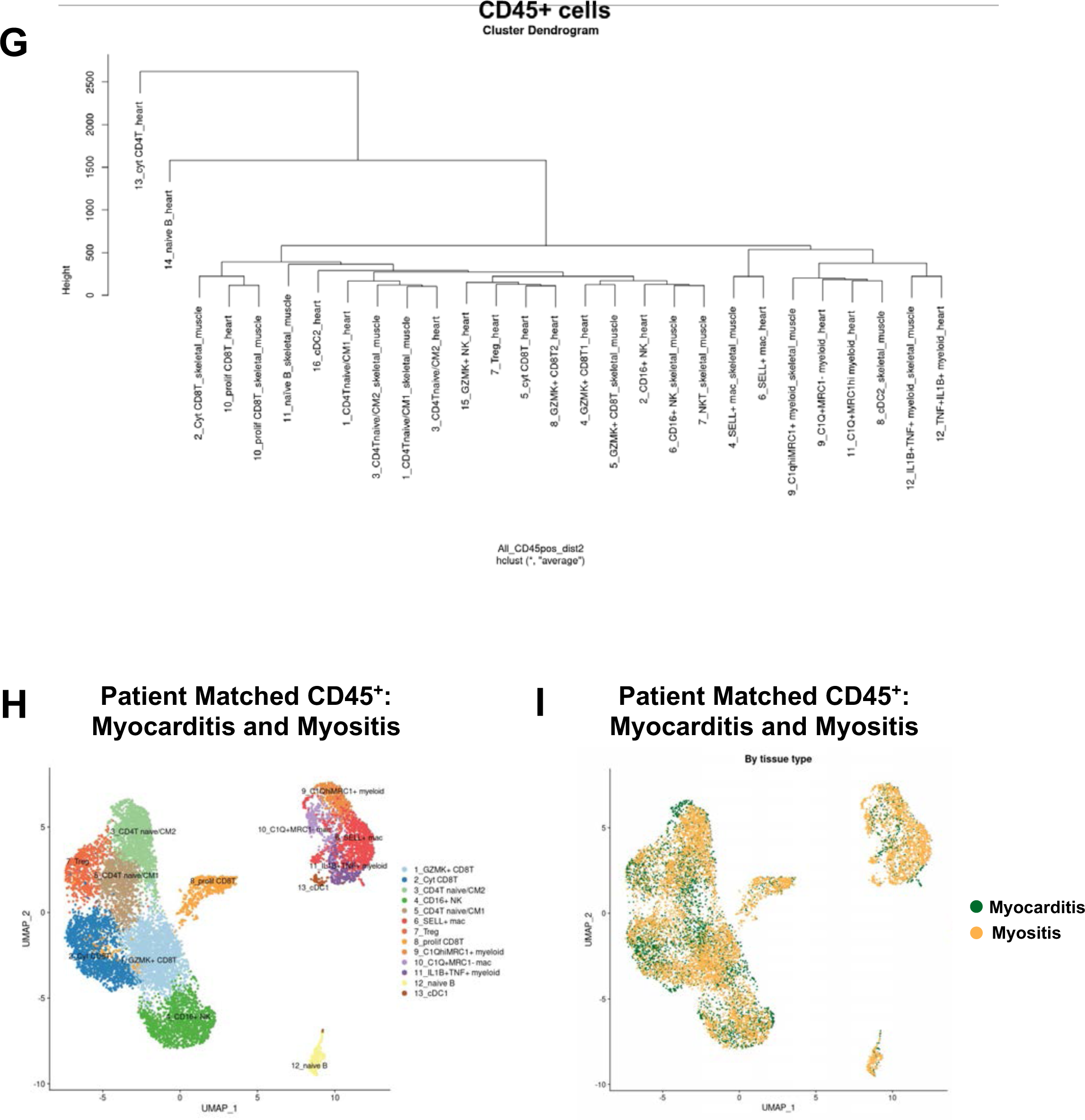
Immune cell subsets in ICT-induced myocarditis and myositis. **A.** UMAP projection of immune cell subclusters in cardiac muscle. **B.** Dot plot indicating expression of individual genes utilized to define subclusters in cardiac muscle. **C.** Comparison of individual cluster frequency in myocarditis versus control samples. **D.** UMAP projection of immune cell subclusters in skeletal muscle. **E.** Dot plot indicating expression of individual genes utilized to define subclusters in skeletal muscle. **F.** Comparison of individual cluster frequency in myositis versus control samples. **G.** Cluster dendrogram of CD45^+^ immune cells in cardiac and skeletal muscle samples. **H.** UMAP projection of patient-matched myocarditis and myositis samples. **I**. Feature plot of patient-matched samples demonstrating relative abundance of cell clusters in myocarditis and myositis samples.

### Clonally-expanded cytotoxic CD8^+^ T cells expressing activation markers are enriched in myocarditis and myositis

Based on prior evidence strongly implicating CD8^+^ T cells in immune-related myocarditis and myositis, we first focused on T cell populations.^3^ In both immune-related myocarditis (**Figures 3a-c**) and myositis **(Figures 3d-f**), we observed a significantly higher frequency of a population of CD8^+^ T cells expressing genes associated with T cell activation (*LAG3, TIGIT, PDCD1, HAVCR2, TOX)* and cytotoxicity (*GZMB, GNLY, NKG7, CST7).* Furthermore, this population of activated, cytotoxic CD8^+^ T cells expressed genes for chemokines, including *CCL5*, *CCL4*, and *CCL4L2* **(Figures 3c & 3e).**^10s" rid="c11">11^ This expression profile is in line with a recent study identifying a circulating population of cytotoxic CD8^+^ T effector cells re-expressing CD45RA (T_EMRA_) cells in the peripheral blood of patients with immune-related myocarditis; these have not previously been directly shown in cardiac tissue from patients.^12^ Within the patient-matched myocarditis and myositis samples, the cytotoxic CD8^+^ T cell subset were clustered similarly (**Figure 3g**) and were present in similar frequency in myocarditis and myositis tissue **(Figure 3h**).

**Figure 3.**
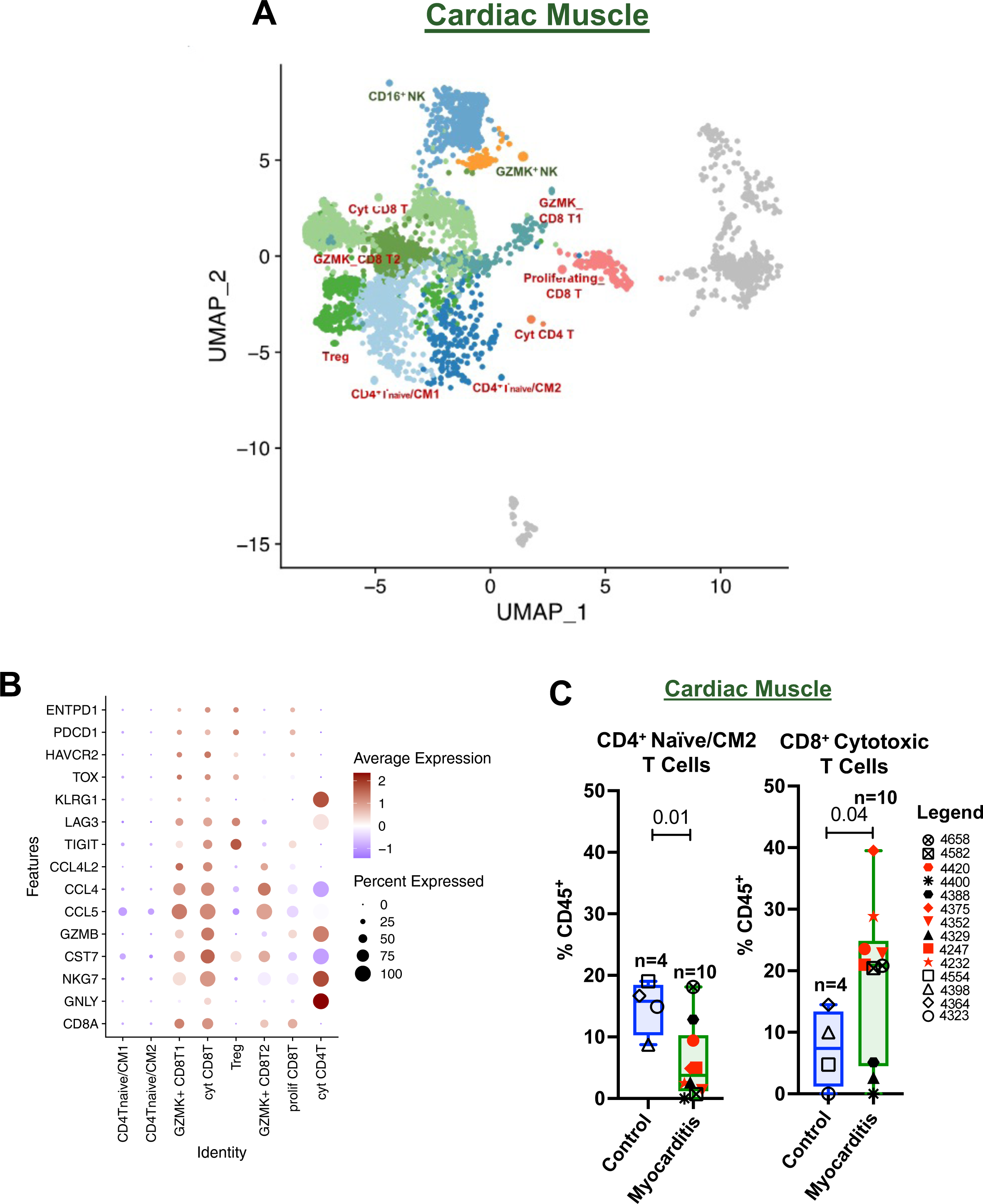
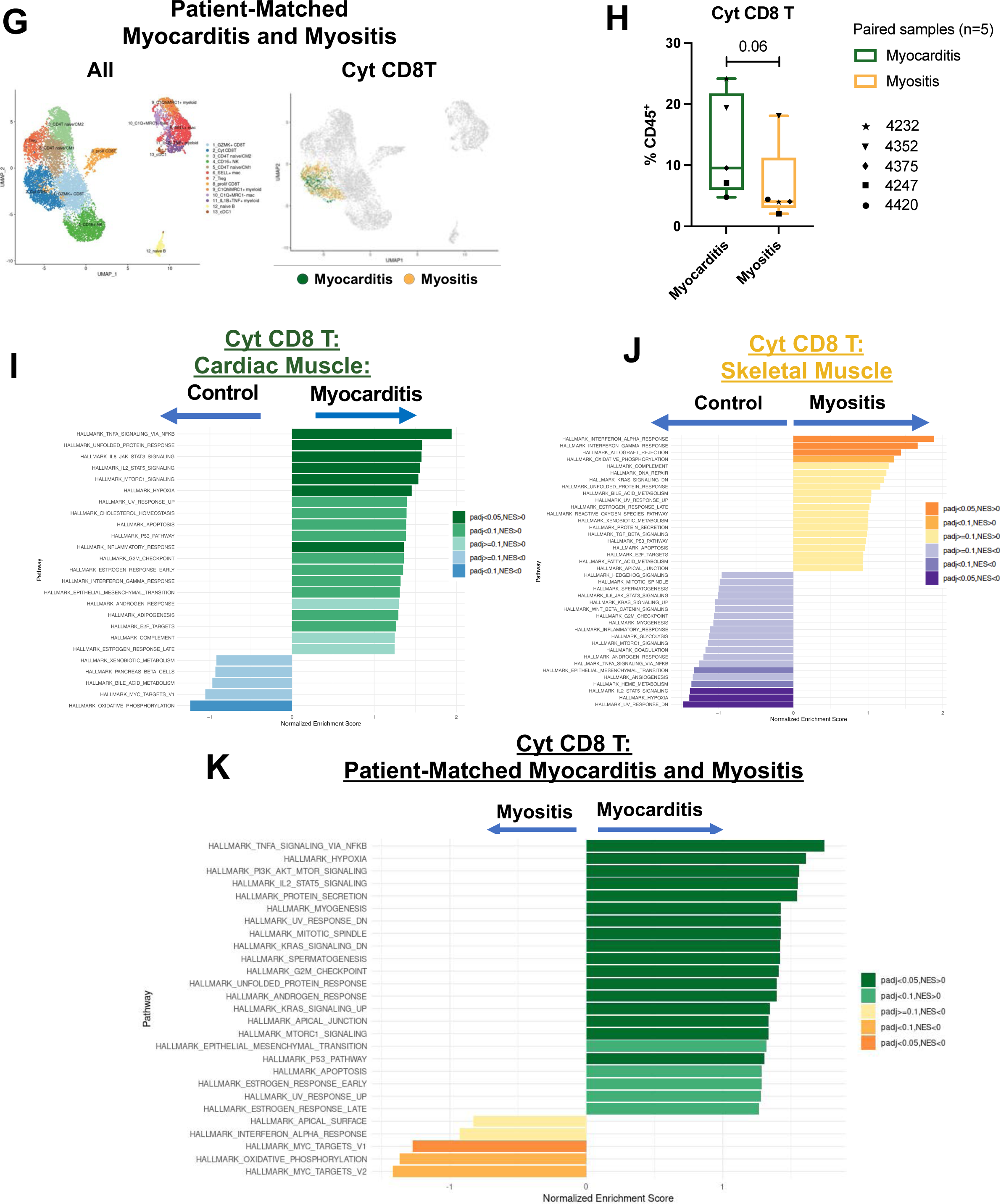
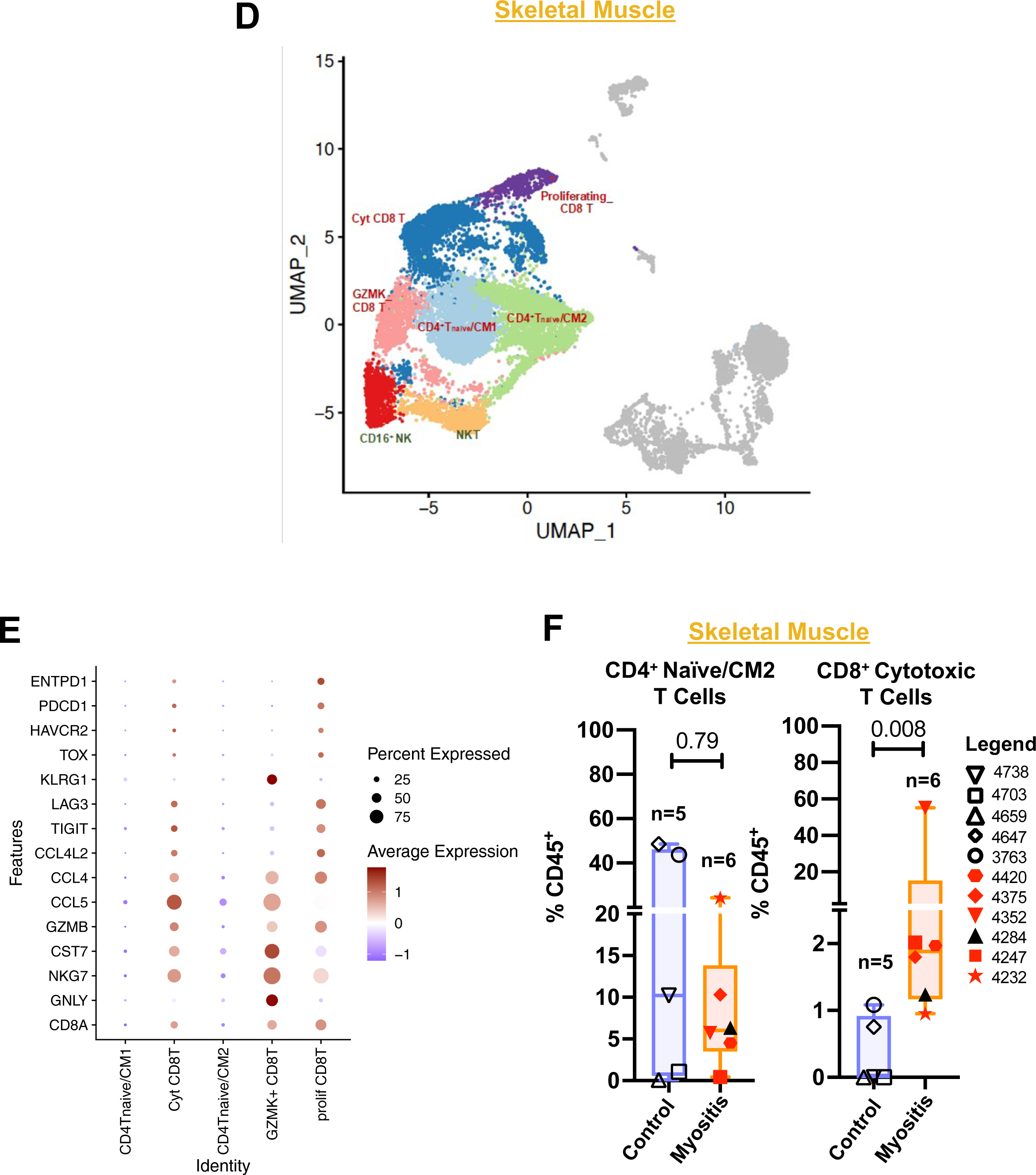
Cytotoxic CD8^+^ T cells expressing activation markers are enriched in myocarditis and myositis. **A.** UMAP projection of individual T cell clusters in cardiac muscle samples. **B.** Dot plot indicating expression of individual genes utilized to define T cell subclusters in cardiac muscle. **C.** Individual T cell subclusters present in differential frequencies in myocarditis versus control cardiac muscle specimens(red indicates patient with matched myocarditis and myositis sample available). **D.** UMAP projection of individual T cell clusters in skeletal muscle samples. **E.** Dot plot indicating expression of individual genes utilized to define T cell subclusters in skeletal muscle. **F.** Individual T cell subclusters present in differential frequencies in myositis versus control skeletal muscle specimens.

As similar marker genes and frequencies were observed within this population in immune-related myocarditis and myositis, we sought to determine if specific transcriptional pathways distinguished these populations in inflamed cardiac and skeletal muscle. The most highly upregulated hallmark transcriptional pathway within the cytotoxic CD8^+^ T cell population in myocarditis compared to control cardiac muscle was TNF-α signaling via NF-κB **(Figure 3i)**, whereas in myositis compared to control skeletal muscle, the most highly upregulated pathway was interferon-α response **(Figure 3j)**. Within the patient-matched specimens, the hallmark hypoxia, IL2/STAT5, and TNF-α signaling via NF-κB signaling pathways were observed to be more highly expressed in myocarditis than in myositis **(Figure 3k)**. In line with this, a subset of differentially expressed genes were shared between the myocarditis and myositis samples, although the transcriptional profiles were not identical, suggestive of context-dependent gene expression within the cardiac and skeletal muscle compartments **(Figure 2g & Supplementary Figures 7a & 7b)**.

Previous studies demonstrated the presence of shared TCR clones between involved heart and skeletal muscles in patients with ICT-induced myocarditis and myositis; however, the specific phenotype of these T cells in inflamed tissue has not previously been described.^2,3^ We therefore analyzed single-cell TCR sequencing (scTCR-seq) data from 9 samples (n=5 myocarditis, n=1 myositis, n=2 control cardiac muscle, and n=1 control skeletal muscle), including one patient with matched myocarditis and myositis samples **(Figure 4 & Supplementary Figure 8)**. For these samples, paired 5’ scRNA-seq data were also generated. We identified a total of 1,477 cells with productive TCR α/β chains after removing cells with multiple TCRs **(Supplementary Figures 8a-b).** Clonotypes were defined based on VJ gene usage and CDR3 nucleotide sequence, and evidence of TCR clonal expansion was observed in toxicity cases **(Supplementary Figures 8c-f)**. Consistent with prior reports, TCR clonal overlap was observed in cardiac and skeletal muscles in the only evaluable patient with matched samples from myocarditis and myositis **(Figure 4a**). We evaluated the specific phenotype associated with these TCR clonotypes, and expanded TCR clones were mapped primarily to a population of cytotoxic CD8^+^ T cells expressing *CCL4* and *CCL5*, consistent with the population we observed to be increased in frequency in myocarditis and myositis tissues **(Figures 4b-d).** Taken together, these findings demonstrate that clonally expanded, cytotoxic CD8^+^ T cells are enriched in cardiac and skeletal muscles of patients with ICT-induced myocarditis and myositis.

**Figure 4.**
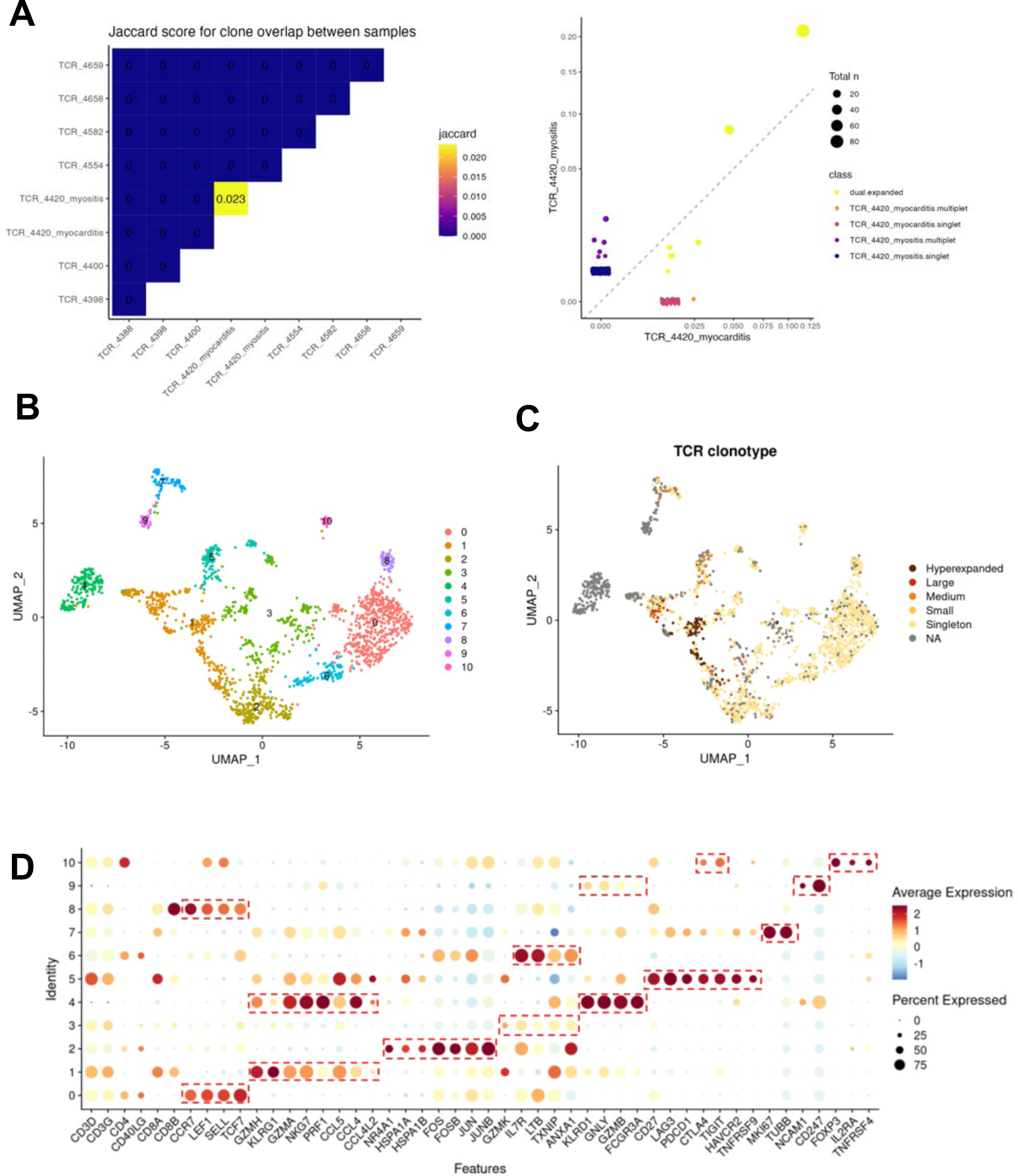
Clonally expanded cytotoxic CD8+ T cells expressing activation markers are shared in myocarditis and myositis. **A.** TCR clonal overlap in matched cardiac and skeletal muscle in patient with myocarditis and myositis. **B.** T and NK cell clusters from the 5’ scRNA-seq dataset. **C.** Mapping of TCR clonotypes to specific T and NK cell clusters. **D.** Marker gene expressions in T and NK cell clusters from the 5’ scRNA-seq dataset.

### Inflammatory IL1B^+^TNF^+^ myeloid cells specifically enriched in ICT-induced myocarditis are associated with poorer clinical outcomes

We next evaluated the myeloid cell compartment, as infiltration of myeloid cells within the inflamed muscle tissue has been reported but their phenotype and putative function have not yet been fully described **(Figures 5a-f & Supplementary Figures 9-10)**.^2,13^ There were no observed differences in the frequencies of tissue-resident C1Q^+^ myeloid populations between the myocarditis and myositis samples compared to control samples **(Figures 5c & 5f).**

**Figure 5.**
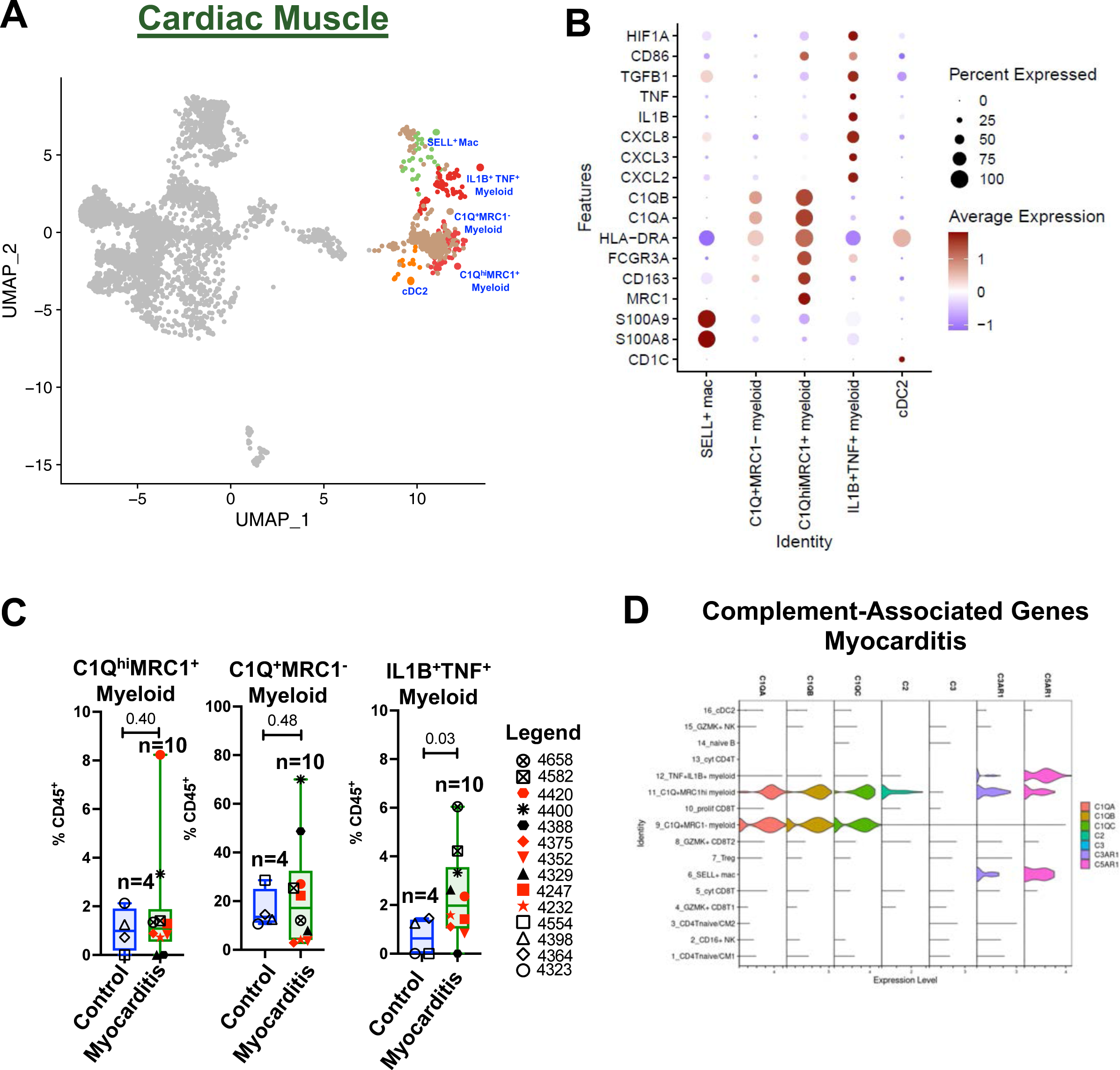
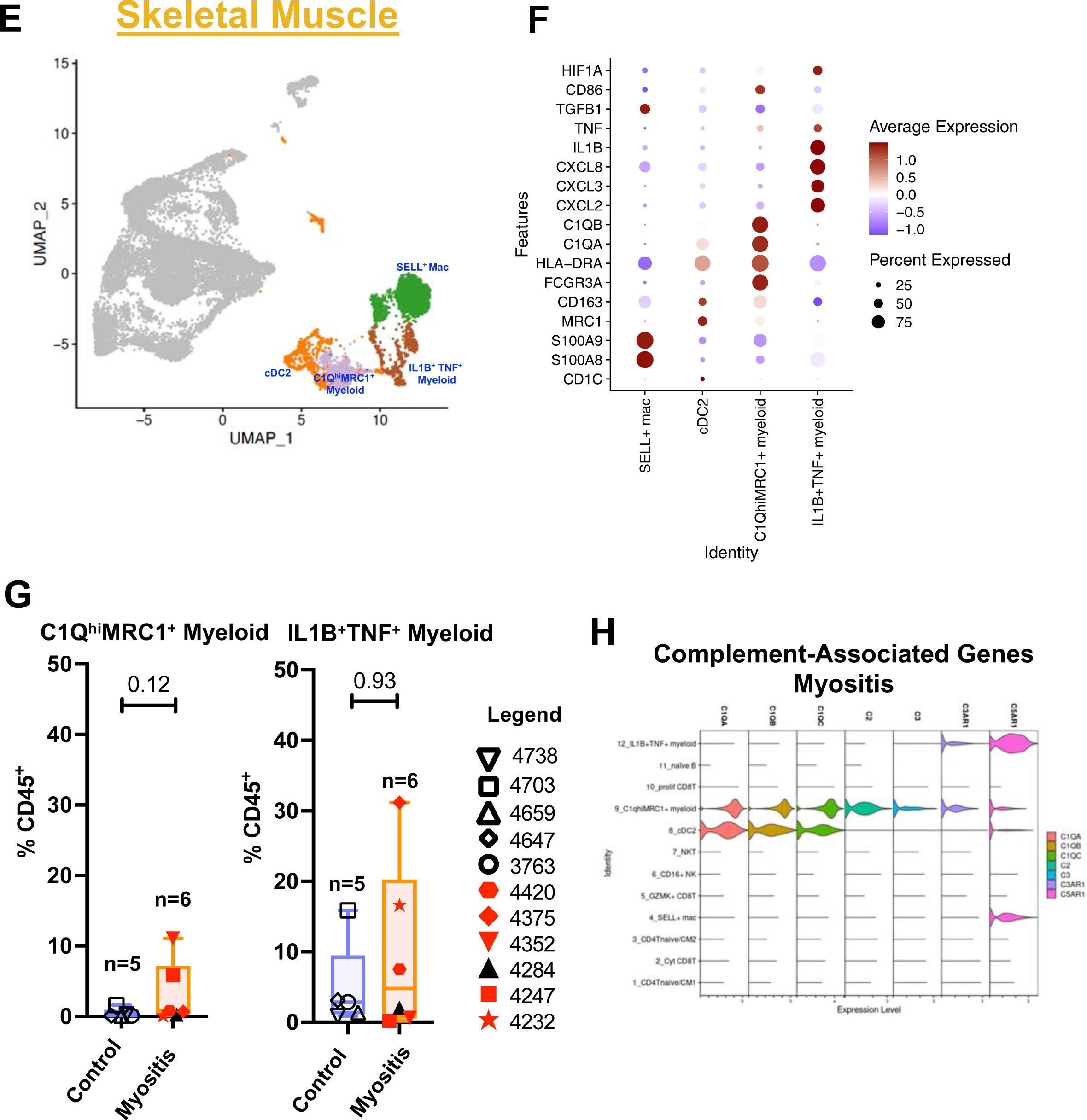
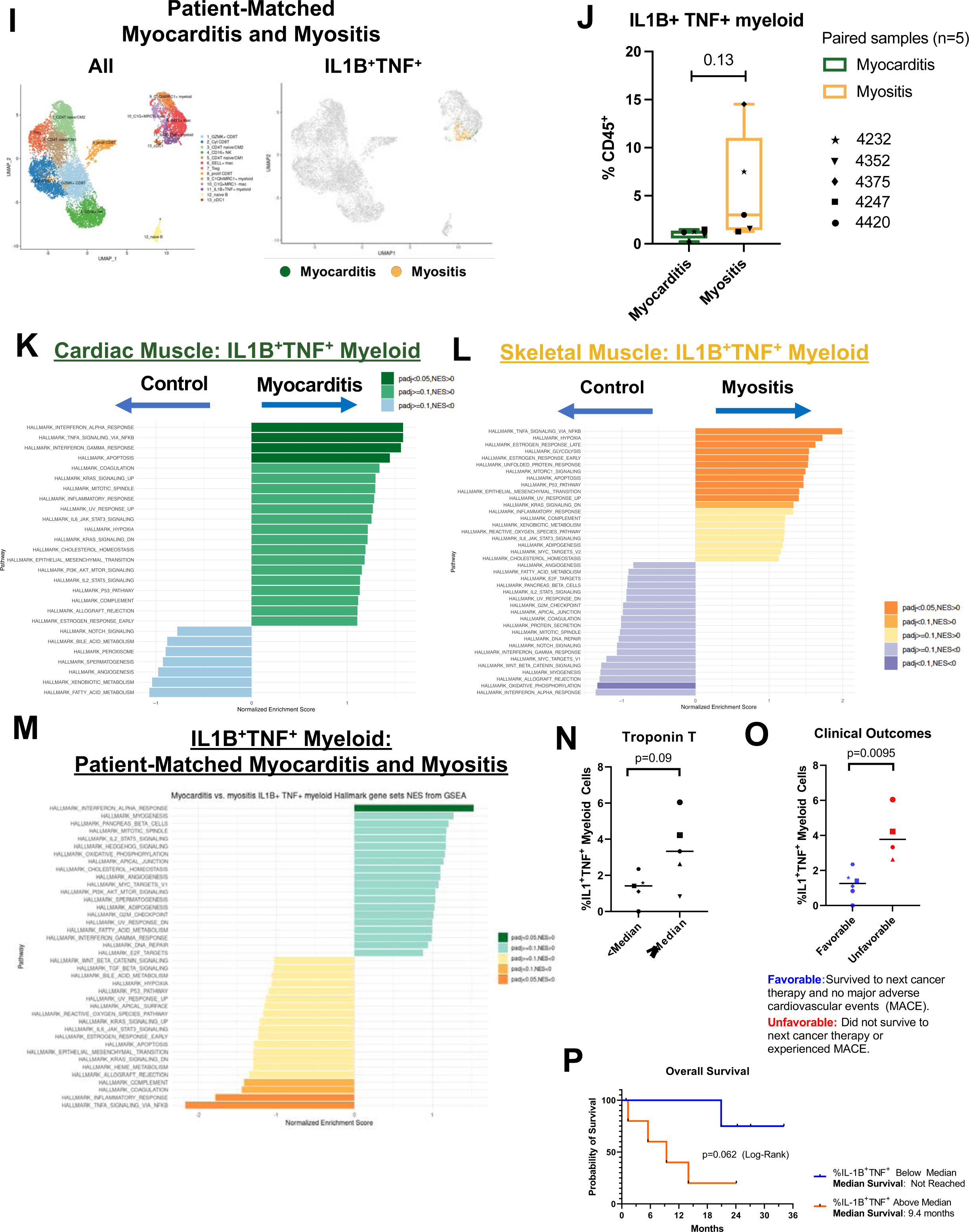
IL-1B^+^TNF^+^ myeloid cells are specifically enriched in ICT-induced myocarditis and are associated with poorer clinical outcomes. **A.** UMAP projection of individual myeloid cell clusters in cardiac muscle samples. **B.** Dot plot indicating expression of individual genes utilized to define myeloid cell subclusters in cardiac muscle. **C.** Frequency of individual myeloid cell subclusters in myocarditis versus control cardiac muscle specimens (red indicates patient with matched myocarditis and myositis sample available). **D.** Violin plot of complement associated gene expression in immune cell subsets in myocarditis samples. **E.** UMAP projection of individual myeloid cell clusters in skeletal muscle samples. **F.** Dot plot indicating expression of individual genes utilized to define myeloid cell subclusters in skeletal muscle. **G.** Frequency of individual myeloid cell subclusters in myositis versus control skeletal muscle specimens (red indicates patient with matched myocarditis and myositis sample available). **H.** Violin plot of complement-associated gene expression in immune cell subsets in myositis samples. **I.** Feature plot of IL1B+TNF+ myeloid cell subclusters in patient-matched myocarditis and myositis samples (right), with all subclusters shown for reference (left). **J.** Frequency of IL1B+TNF+ myeloid cell subpopulations in patient-matched myocarditis versus myositis samples. **K.** Differentially upregulated transcriptional pathways within IL1B+TNF+ myeloid cell population in myocarditis versus control cardiac muscle. **L.** Differentially upregulated transcriptional pathways within IL1B+TNF+ myeloid cell population in myositis versus control skeletal muscle. **M.** Differentially upregulated transcriptional pathways within IL1B+TNF+ myeloid cell population in patient-matched myocarditis vs. myositis. **N.** Association of frequency of IL1B+TNF+ myeloid cells with serum troponin T. **O.** Association of frequency of IL1B+TNF+ myeloid cells with clinical outcomes. **P.** Overall survival stratified by frequency of IL1B+TNF+ myeloid cells.

Of note, the population of tissue-resident C1Q^hi^MRC1^+^ myeloid population expressed *FCGR3A* **(Figures 5b & 5f and Supplementary Figures 11a-c)**, the gene for Fc_γ_RIIIa, a receptor that recognizes IgG antibodies and regulates complement activation, a potential mediator of tissue injury. Binding of IgGs to specific Fc receptors (including Fc_γ_RIIIa) leads to activation of the classical complement pathway and induction of antibody-dependent cellular cytotoxicity (ADCC). Consistent with a potential role for these cell types in complement activation, we observed upregulation of classical complement pathway genes, including *C1QA, C1QB, C1QC*, and *C2* within the population of tissue-resident C1q^hi^MRC1^+^ myeloid cells **(Figures 5d & 5i and Supplementary Figure 11d).** To confirm these findings at the protein expression level, we performed immunohistochemistry (IHC) on 13 formalin-fixed paraffin-embedded tissues evaluable for further analysis (n=6 myocarditis; n=3 myositis; n=3 control cardiac; n=1 control skeletal) **(Supplementary Figure 11e).** Consistent with our scRNA-seq findings, we observed infiltrating T cells (CD8^+^ > CD4^+^) and CD68^+^ myeloid cells (macrophages) within inflamed cardiac and skeletal muscle tissues (**Supplementary Figure 11e).** We identified protein expression of the complement split product C4d and pan-IgG on all evaluable immune-related myocarditis and myositis samples tested (**Supplementary Figures 11e & f)**.^14^ These findings were associated with detectable serum autoantibodies (anti-striated muscle or anti-acetylcholine receptor binding or modulating IgGs) in 55% of our patients with confirmed toxicity (6/11) **(Supplementary Figure 11g & Supplementary Table 1).** Notably, positive serum striated muscle or acetylcholine receptor IgGs were observed in 50% of myocarditis specimens (3/6) and 33% of myositis specimens (1/3) **(Supplementary Figure 11c),** suggesting the presence of other as-yet uncharacterized autoantibodies in immune-related myocarditis and myositis.

We further observed a significantly higher frequency (p=0.03) of HLA-DRA^lo^CD86^+^ myeloid cells expressing the pro-inflammatory cytokines *IL1B* and *TNF*, the pleiotropic cytokine *TGFB1*, and the hypoxia-response factor *HIF1A* within the immune-related myocarditis samples **(Figures 5b & 5c)**. This population expressed *CXCL2* (also called macrophage inflammatory protein [MIP]-2α, chemotactic for granulocytes and hematopoetic stem cells), *CXCL3* (MIP-2β, which is chemotactic for monocytes), and *CXCL8* (IL-8, mediates neutrophil chemotaxis) **(Figure 5b).** This population was also identified within skeletal muscle tissues but was not significantly increased (p=0.93) in myositis compared to control samples **(Figures 5f & 5g)**. Quantitative comparison of the transcriptomes of these populations between myocarditis and myositis samples demonstrated that a plurality of differentially expressed genes were shared **(Supplementary Figure 12).** Furthermore, within the patient-matched myocarditis and myositis samples, the IL1B^+^TNF^+^ myeloid cell subset were clustered similarly (**Figure 5i**) and were present in comparable frequencies in myocarditis and myositis tissues **(Figure 5j**).

We sought to determine if specific transcriptional pathways distinguished these myeloid cell populations in inflamed cardiac and skeletal muscle. Within cardiac muscle, the most highly expressed transcriptional pathways in this population of myeloid cells were IFN-α response, TNF-α signaling via NF-κB, and IFN-γ response **(Figure 5k)**. In skeletal muscle, TNF-α signaling via NF-κB was the most highly expressed transcriptional pathway, however the IFN-α response and IFN-γ response pathways were not significantly upregulated in myositis **(Figure 5l)**. Consistent with this, direct comparison of matched myocarditis and myositis samples identified significantly increased interferon-α response pathway in myocarditis and significantly increased TNF-α signaling via NF-κB in myositis **(Figure 5m).**

As this population of IL1B^+^TNF^+^ myeloid cells has not previously been identified in these toxicities and was found to be significantly increased in myocarditis, we evaluated the association with severity of myocarditis and clinical outcomes. We observed a trend toward higher median serum troponin-T levels associating with increased frequency of the IL1B^+^TNF^+^ myeloid cell subset (p=0.09) **(Figure 5n)**. Furthermore, a higher frequency of the IL1B^+^TNF^+^ myeloid cell subset was associated with poorer clinical outcomes, defined as an occurrence of a major adverse cardiovascular event (a composite of heart failure, arterial thrombus, life threatening arrhythmia, pulmonary embolism or unexplained sudden cardiac death) or death before next anti-cancer therapy, whereas a higher frequency of the cytotoxic CD8^+^ T cell population was not associated with poorer clinical outcomes **(Figure 5o & Supplementary Figure 12c).** Moreover, we observed that an increased frequency of IL1B^+^TNF^+^ myeloid cells (above the median) was associated with a trend toward shorter overall survival (p=0.062) **(Figure 5p).** The duration of survival is consistent with reports from our group and others that non-fatal irAEs are associated with improved overall outcomes following ICTs.^15–17^ To evaluate potential interactions between these myeloid cells, T cells, and stromal cells, we performed computational cell-cell communication analyses and identified predicted interactions of cytotoxic CD8^+^ T cell-derived IFN-γ and IL1B^+^TNF^+^ myeloid cell-derived IL1β on endothelial cells, consistent with endothelial stress responses **(Supplemental Items 1-5).**

## Discussion

ICTs offer the promise of long-term survival benefit to subsets of patients with cancer, but can induce life-threatening irAEs. We focused here on immune-related myocarditis and myositis, rare irAEs affecting muscle tissues that carry high mortality. There is an unmet need to understand the molecular pathways and immune subsets underlying these irAEs to inform rational, front-line immunosuppressive treatments. We performed single-cell analyses of cardiac and skeletal muscle tissues at the time of hospitalization since performing baseline endomyocardial and quadriceps muscle biopsies in otherwise healthy patients represented an unacceptable risk. The strength of our approach is that these technically challenging biopsies were obtained early in the course of toxicity in living patients, before progression to mortality, which may lead to cellular breakdown and confound evaluation of the inflamed microenvironment. Furthermore, all patients except for one had overlapping myocarditis and myositis, and we did not observe a skew in our dataset from the one patient with myocarditis-only. It should be noted that clinically necessary treatments could not be withheld prior to biopsy. Therefore, most patients received at least one dose of intravenous corticosteroids, the first-line treatment for these irAEs, potentially limiting yield of CD45^+^ cells in these needle biopsies. Despite these limitations, we identified molecular pathways and immune subsets in myocarditis and myositis that may inform investigation into rational treatments for these irAEs.

Our data identified the enrichment of clonally-expanded cytotoxic CD8^+^ T cells in both myocarditis and myositis and inflammatory myeloid cells expressing genes for the TNF-α and IL-1β cytokines specifically in immune-related myocarditis. It should be noted that this population of myeloid cells was also identified in myositis but was not significantly different in frequency between myositis and control samples, possibly due to the higher cellular frequency observed in both groups. TNF-α promotes further release of inflammatory cytokines, upregulation of endothelial adhesion molecules, and regulation of leukocyte migration.^18–20^ In fact, TNF inhibitors have successfully been utilized as second-line treatment of immune-related myocarditis; however, their clinical applicability is limited by the increased risk of infection and congestive heart failure (CHF), particularly at higher doses.^21,22^ Another second-line approach is blockade of the T cell CD28-CD80/86 costimulatory pathway with the CTLA-4 Ig fusion protein, abatacept, which has been shown to provide clinical benefit in a case series of patients with immune-related myocarditis alone and in combination with Janus kinase 1 (JAK1) and JAK2 inhibitor, ruxolitinib.^23,24^ The identification of upregulated IL6/JAK/STAT3 signaling in our data supports this role of ruxolitinib. Furthermore, our data suggest that abatacept may function through disruption of costimulatory signaling via CD86 provided by the identified population of inflammatory IL-1B^+^TNF^+^ myeloid cells.

Myeloid cells have been implicated in the development of irAEs, such as colitis and arthritis, and have heterogeneous effects on anti-tumor immunity.^4,5,25^ Therefore, it is important to understand the relative contributions of myeloid subpopulations to toxicities and anti-tumor immune responses. Our study is the first to identify IL-1B-expressing myeloid cells associated with poorer clinical outcomes in immune-related myocarditis, suggesting that the IL-1 pathway is an attractive target for selective immunosuppression. IL-1 encompasses two cytokines: IL-1α, which primarily functions in local inflammatory responses, and IL-1β, which is produced chiefly by myeloid cells (including macrophages, neutrophils and mast cells) and exerts systemic effects. Engagement of the IL-1 receptor initiates multiple downstream events, including production of inflammatory cytokines such as IL-6, induction of fever, promotion of leukocyte migration, and supporting expansion and effector functions of CD4^+^ and CD8^+^ T cells.^26–28^ Indeed, in a murine model of ICT-induced myocarditis, IL-1β blockade was found to prevent myocarditis and reduce cardiac fibrosis.^29,30^ However, it should also be noted that IL-1 signaling has not been found to be consistently upregulated in all murine models of myocarditis.^3^ Furthermore, the role of inflammatory IL-1 signaling in non-ICT-related heart disease, including viral myocarditis and coronary artery disease, has also previously been recognized.^31–33,34,35^ In fact, IL-1 pathway inhibitors are currently in clinical use for the treatment of pericarditis, inflammation of the lining of the heart,^36^ and an anti-IL-1β antibody (canakinumab) was shown to reduce cardiovascular events in patients with prior myocardial infarction.^31–33^ A clinical trial targeting the IL-1 pathway in patients immune-related myocarditis is currently under development at our institution.

Our data further implicate B cell responses, with secretion of IgGs that recognize muscle antigens potentially engaging with Fc_γ_RIIIa on tissue-resident C1q^hi^MRC1^+^ myeloid cells leading to downstream activation of the complement pathway, as observed both transcriptionally and at the protein level, with deposition of the complement split product C4d. These findings provide a rationale for B cell-directed therapies such as plasma exchange (to remove autoantibodies) and anti-CD20 antibodies (to deplete B cells) in myocarditis, myositis, and other associated irAEs such as myasthenia gravis, Guillain-Barre syndrome (GBS), vasculitis, and glomerulonephritis and may suggest a role for direct complement-targeting therapies in these toxicities. This potential role for B cells in irAEs is aligned with recent reports of B cells organized into tertiary lymphoid structures (TLS) contributing to anti-tumor immune responses with ICTs.^37–39^

In summary, we identified populations of clonally expanded cytotoxic CD8^+^ T cells expressing activation markers shared in cardiac and skeletal muscle and specific myeloid cell subsets that may be involved in immune-related toxicities via the IL-1 and classical complement pathways. These results nominate candidate clinically actionable targets for investigation in these severe irAEs, to ultimately advance the complementary goals of minimizing toxicity and maximizing efficacy of ICTs.

## Methods

### Study Design and Patients

Patients hospitalized at The University of Texas MD Anderson Cancer Center from November 2020 to February 2022 with suspected immune-related myocarditis, myositis, and myasthenia gravis were consented to an institutional review board (IRB)-approved laboratory protocol (MD Anderson PA13-0291) for immune monitoring studies. All patients underwent standard diagnostic testing and treatment as clinically indicated. Patients with suspected immune-related myocarditis underwent endomyocardial biopsy by an interventional cardiologist with at least four cores sampled. Patients with suspected immune-related myositis underwent skeletal muscle (quadriceps) biopsy by an interventional radiologist. Cardiac and skeletal muscle pathology specimens were reviewed by a clinical pathologist to assign a clinicopathologic diagnosis. ^7,8^ All biopsies were obtained within 96 hours of clinical presentation. Due to the life-threatening nature of these toxicities, corticosteroids were not withheld prior to biopsy. In most cases, biopsies were obtained within 24 hours of corticosteroid initiation. Confirmed irAE diagnosis required all three of the following criteria: 1. symptoms requiring hospitalization; 2. elevated serum markers (creatine kinase [CK, muscle enzyme] ≥5x upper limit of normal, troponin-I or troponin-T [cardiac enzyme] >99th percentile or presence of autoantibodies known to be associated with MG); 3. positive muscle tissue biopsy, electromyography (EMG), or cardiac magnetic resonance imaging (MRI).

### Single-Cell RNA Sequencing

scRNA-seq was performed using the 10x Genomics Chromium Single Cell Controller. Briefly, single-cell suspensions were prepared from cardiac and skeletal muscle biopsies as described.^13^ Cells were resuspended in freezing media containing 90% AB serum (derived from donors with AB blood type) and 10% dimethyl sulfoxide (DMSO) and stored in liquid nitrogen until analysis. For scRNA-seq analysis, cells were thawed, washed, and droplet-separated using the Chromium Single Cell 3′ v.3 Reagent Kit (10x Genomics) or 5′ v.2 Reagent Kit (10X Genomics) with the 10x Genomics microfluidic system creating cDNA library with individual barcodes for individual cells. Barcoded cDNA transcripts from patients were pooled and sequenced using the NovaSeq 6000 Sequencing System (Illumina). For single-cell TCR sequencing, an aliquot of cDNA generated by 5′ v.2 Reagent Kit was used to enrich for V(D)J regions using the Chromium Single Cell V(D)J Enrichment Kit Human T Cell (10X Genomics) according to the manufacturer’s protocol. V(D)J region-enriched libraries were pooled and sequenced using the NovaSeq 6000 Sequencing System (Illumina).

### Overview of 3’ Single-Cell RNA sequencing analysis

Raw scRNA-seq reads generated by Illumina sequencer were demultiplexed into FASTQ and aligned to GRCh38 reference genome to generate count matrices using Cell Ranger v6.0.0 analysis pipelines (10x Genomics). The preliminarily processed data were used for downstream analysis by Seurat v4.3.0.^44^ Low-quality and dead cells which had <200 or >>2,500 unique transcripts or >5% of mitochondrial transcripts were filtered out. Multiple datasets (control, myocarditis, and myositis) were combined, normalized, identified variable features, and applied principal component analysis (PCA) using Seurat. Batch effect correction of combined data was performed using Harmony algorithm v0.1.1.^45^ Following batch correction, we applied FindClusters with a resolution of 0.5 to identify cell clusters and further non-linear dimension reduction was performed with uniform manifold approximation and projection (UMAP).

### Comparison of transcriptomic profiles among cell types in myocarditis and myositis samples

Overlapping DEGs from the two datasets (myocarditis and myositis) were identified within the same cell type. Overrepresentation analysis was then performed on these overlapping DEGs against the hallmark gene sets in the Molecular Signature Database^46–48^ using the R package clusterProfiler (version 4.4.4).^49^ Briefly, the analysis of overrepresented pathways was performed with the function *enricher*. Then, the results were visualized with the function *dotplot* in clusterProfiler.^49^

### 5’ scRNA-seq data analysis

#### Raw data preprocessing and quality control

The raw sequencing data were preprocessed (demultiplexing cell barcodes, read alignment, and generation of gene count matrix) using the Cell Ranger Single Cell Software Suite (version 7.1.0) provided by 10× Genomics. The human reference genome GRCh38 was used for sequence alignment. Subsequent analysis was performed using the Seurat software (version 4.3.0).^44^ For quality filtering, genes detected in fewer than 10 cells and cells with transcripts aligned to less than 200 genes were removed. Cells with 15% or more of their transcripts aligned to mitochondrial genes were also filtered out. Furthermore, cells with transcripts aligned to 6,000 or more genes were removed to exclude potential doublets or multiplets. Following these steps, a total of 10,861 cells were retained for downstream analysis. Following this initial clustering, we removed likely doublets and multiplets, as we previously described.^50^

#### Library size normalization, dimension reduction, batch correction and unsupervised clustering

Log normalization on the library size of each cell was applied to the gene count matrix. 2,000 highly variable genes were identified with the function *FindVariableFeatures* in Seurat.^44^ The data were then scaled, and a linear dimension reduction based on these highly variable genes was conducted using the *RunPCA* function in Seurat.^44^ An elbow plot was drawn, and the number of principle components used for subsequent analysis was determined accordingly. Batch effects were evaluated, and batch correction was conducted with Harmony (version 0.1.1) software in R.^45^ A nearest-neighbor graph was constructed using the *FindNeighbors* function in Seurat.^44^Following this, unsupervised clustering was performed with the Louvain algorithm at a resolution of 0.1, resulting in the identification of 12 cell clusters. Non-linear dimension reduction was performed using the Uniform Manifold Approximation and Projection (UMAP) method. Differential gene expression analysis was then conducted with the *FindAllMarkers* function in Seurat,^44^ using the Wilcoxon Rank Sum test, with the *min.pct* parameter set to 0.25. Major cell types were further determined based on a manual review of DEGs and the expression of classical lineage markers in each cluster. By checking multiple lineage markers, we observed no additional doublets or multiplets at the major cell type level.

#### Sub-clustering analysis of T & NK cells

We further performed sub-clustering on T & NK cells, following the same procedures as described above. Cell clusters that simultaneously expressing discrepant lineage makers (e.g., cells in the T-cell cluster showed expression of B cell markers) were identified and removed. We performed several rounds of filtering as we previously reported until no obvious doublets or multiplets were observed.^50^ DEGs were identified with the same steps as above. Top 30 most significant DEGs were carefully reviewed. Besides, the expressions of canonical markers for T cell types and states, as reported previously,^51^ were manually checked, and bubble plots were generated accordingly. T cell types and states were then inferred by integrating the above information, and annotations were added to each cluster with the *AddMetaData* function in Seurat. ^44^

#### Assessing the transcriptome similarity among defined cell states

To assess the transcriptome similarity and phenotypic relationships among the T cell states identified in this study, the CD45^+^ cells from the 3’ and 5’ scRNA-seq datasets were merged, followed by library size normalization, dimension reduction, batch correction and unsupervised clustering analysis as described above. The distribution of the cells from the 3’ and 5’ scRNA-seq datasets in the UMAP space was visualized using the R package ggplot2 (version 3.4.0) (Wickham, 2016) and the *DimPlot* function in Seurat.^44^ In addition, we performed a hierarchical clustering analysis. Briefly, the Harmony embeddings were first extracted with the *Embeddings* function in Seurat.^44^ Then, Euclidian distances between cell clusters were calculated based on the average Harmony space for each cell cluster, and a dendrogram was drawn based on hierarchical clustering to show the relationships of cell types defined in the 3’ and 5’ scRNA-seq datasets.

### scTCR-seq data analysis

The preprocessing of raw sequencing data was conducted using the Cell Ranger software (version 7.1.0, 10× Genomics) with the *cellranger vdj* function. The pipeline encompassed several steps, including cell barcode demultiplexing, aligning reads to the human reference genome GRCh38, assembling TCRs, and identifying the V(D)J gene usage and sequences of the framework regions and complementary determining regions (CDRs). For downstream analysis, the R package scRepertoire (version 1.7.0) was used.^52^ As part of the quality control, only productive TCRα and TCRβ chains were kept, and cells with multiple TCRα or TCRβ chains were removed. TCRs with exactly the same VJ gene usage and CDR3 sequences on both TCRα and TCRβ chains were assigned as the same clonotype. The clonal size of TCRs was classified as hyperexpanded (clonal proportion > 0.10), large (0.05 < clonal proportion ≤ 0.10), medium (0.01 < clonal proportion ≤ 0.05), small (clonal proportion ≤ 0.01) and singleton (with only 1 cell expressing the clonotype).

The inverse Simpson index was used to assess clonal diversity. It is calculated as follows:

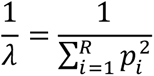

where 𝜆 represents the Simpson index, 𝑝_𝑖_ represents the proportion of clonotype 𝑖 in the repertoire, and 𝑅 represents the total number of clonotypes.

To assess the level of clonal overlap between samples, the Jaccard index was employed, which is calculated as follows:

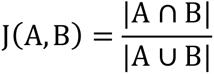

where A and B represents the two samples being compared; J(A, B) denotes the Jaccard index between A and B. A ∩ B represents the size of the intersection between A and B, and A ∪ B represents the size of the union of A and B. Heatmaps were used to visualize the Jaccard scores between samples. In addition, alluvial plots were employed to illustrate the overlap of the top TCR clones across different samples.

The TCR clonotype information was further integrated with the 5’ scRNA-seq data based on the unique cell barcodes, using the *AddMetaData* function in Seurat.^44^ The clonal size distribution in each T cell cluster was visualized using the R package ggplot2 (version 3.4.0).

### Cell-cell communication analysis

Cell-cell communication analysis was conducted using the NicheNet method, which utilizes the information of active target signaling pathways to infer cellular interactions. ^53^ The analysis was carried out using the nichenetr package (version 1.1.1) in R. Cellular interactions in different conditions (myocarditis vs. normal myocardium, myositis vs. normal skeletal muscle) were compared using the *nichenet_seuratobj_aggregate* function. Additionally, the comparison of cellular interactions across different target cell types was performed with the *nichenet_seuratobj_cluster_de* function, also part of the nichenetr package. For the identification of active ligands, only upregulated target genes in the condition or cell type of interest were taken into consideration.

### Serum Autoantibodies

IgGs were evaluated in patient serum at the time of hospitalization using Clinical Laboratory Improvement Amendments (CLIA)-certified striated muscle and myasthenia gravis/Lambert-Eaton myasthenia syndrome antibody panels, including autoantibodies for acetylcholine receptor (muscle) binding antibody with reflex testing to acetylcholine receptor modulating antibody.

### Immunohistochemistry

At least three samples from the right ventricular endomyocardial biopsy and quadriceps biopsy were fixed in phosphate-buffered formalin and embedded in a single paraffin block. Sections were then made for evaluation with light microscopy including staining for hematoxylin and eosin (H&E), trichrome and immunohistochemistry (IHC). IHC staining was performed for CD4 (Agilent IR649, Clone 4B12), CD8 (Agilent GA623, Clone C8/144B), CD20 (Agilent GA604, Clone L26), CD68 (Agilent GA609, Clone KP1), pan-IgG (Agilent IR512, polyclonal), and C4d (Biorad 0300-0230, polyclonal). One sample from right ventricular endomyocardial biopsy was fixed in glutaralde-hyde and electron microscopy was performed (results not presented).

### Statistical analysis

Normality assumption of data was assessed using Shapiro-Wilk test. For datasets with normal distributions (p > 0.05), a two-tailed Welch’s *t*-test was employed. For datasets with non-normal distributions (p < 0.05), a two-tailed Mann-Whitney *U*-test was used. For the comparison of cell proportions in the patient-matched dataset, Wilcoxon matched-pairs signed rank tests were performed. Statistical tests and data visualizations were performed using Prism 9.4.1 (GraphPad Software) and R statistical software (version 4.2.0).”

## Declarations

### Acknowledgements

We are grateful to our patients and their families for their participation in this research. We thank the Immunotherapy Platform at MD Anderson for collecting and processing tissue biopsies and generating scRNA-seq gene expression libraries, and the Advanced Technology Genomics Core at MD Anderson for performing sequencing.

### Authors’ contributions

The first draft of the manuscript was written by B.A.S. All authors contributed to the final manuscript and approved the decision to submit the manuscript for publication. The corresponding authors (S.K.S. and P.S.) had access to all data and had final responsibility for the decision to submit for publication. S.B. and Z.H. designed scRNA-seq experiments. L.Y supervised bioinformatics data analysis approaches. Z.H. and Y.D. performed computational analysis. S.B., Z.H., B.A.S, N.L.P, L.Y, S.K.S. and P.S. interpreted the data

### Funding

These studies were supported by the Parker Institute for Cancer Immunotherapy.

### Competing Interests

B.A.S., S.B., Z.H., M.B., C.A.I., A.D., A.R-K., S.S.Y., and R.S. report no competing interests. N.L.P. reports support by Cancer Prevention & Research Institute of Texas Early Clinical Investigator Award, NIH/NCI, and the Andrew Sabin Family Foundation; consulting relationships with Replimmune, Kiniksa, and honoraria from Physician Education Resource, Society for Immunotherapy for Cancer, and Scripps. L.M.B reports honorarium from Elsevier and Springer. J.P.A is a Scientific Advisory Committee Member for Achelois, Affini-T, Apricity, Asher Bio, BioAtla LLC, Candel Therapeutics, Catalio, Carisma, Codiak Biosciences, Inc, C-Reveal Therapueutics, Dragonfly Therapeutics, Earli Inc, Enable Medicine, Glympse, Henlius/Hengenix, Hummingbird, ImaginAb, Infinity Pharma, InterVenn Biosciences, LAVA Therapeuticss, Oncolytics, PBM Capital, Phenomic AI, Time Bioventures, Two Bear Capital, and Xilis,Inc. He has private investments with Adaptive Biotechnologies, BioNTech, JSL Health, Sporos, and Time Bioventures. P.S is a Scientific Advisory Committee Member for Achelois, Adaptive Biotechnologies, Affini-T, Apricity, Asher Bio, BioAtla LLC, Candel Therapeutics, Catalio, Carisma, Codiak Biosciences, Inc, C-Reveal Therapueutics, Dragonfly Thereapeutics, Earli Inc, Enable Medicine, Glympse, Henlius/Hengenix, ImaginAb, Infinity Pharma, InterVenn Biosciences, JSL Health, LAVA Therapeuticss, Oncolytics, PBM Capital, Phenomic AI, Time Bioventures, Two Bear Capital, and Xilis,Inc. She has private investments with BioNTech, JSL Health, Sporos, and Time Bioventures. S.K.S. reports Consulting or Advisory Role: Amgen, Apricity Health LLC, Arcus Biosciences, Bayer, Boxer Capital, Breaking Data, Bristol-Myers Squibb, Cancer Expert Now, ChemoCentryx, Dendreon, InProTher, Kahr Medical Ltd, Janssen, Javelin Oncology, MD Education Limited, Merck, OncLive (Owned by Intellisphere, LLC), Pfizer, Portage, Regeneron and The Clinical Comms Group; Research.

### Availability of data and material

Data are available upon reasonable request. All data relevant to the study are included in the article or uploaded as supplementary material.

---

**Supplementary Figure 1:**
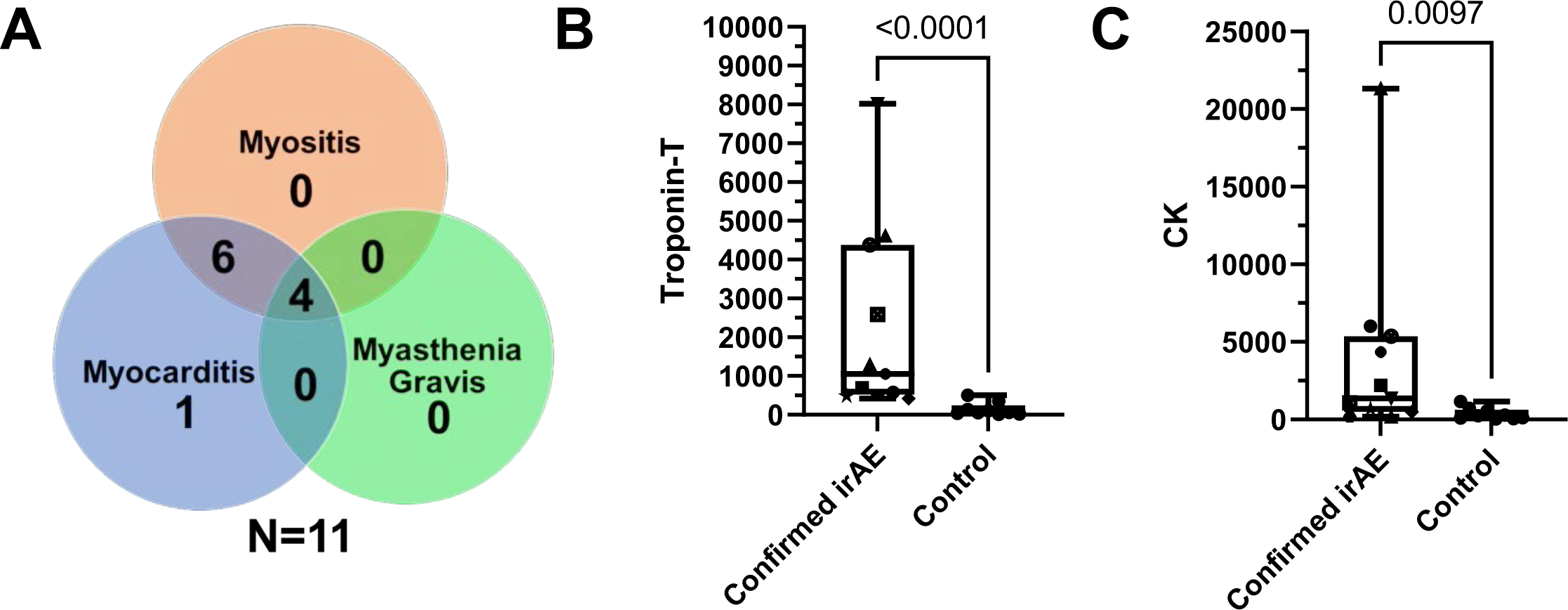
Clinical characteristics of patients with confirmed toxicities. **A.** Distribution of irAE cases. **B**. Serum troponin T measurements in confirmed irAE cases compared to controls. **C.** Serum creatine kinase (CK) measurements in confirmed irAE cases compared to controls.

**Supplementary Figure 2:**
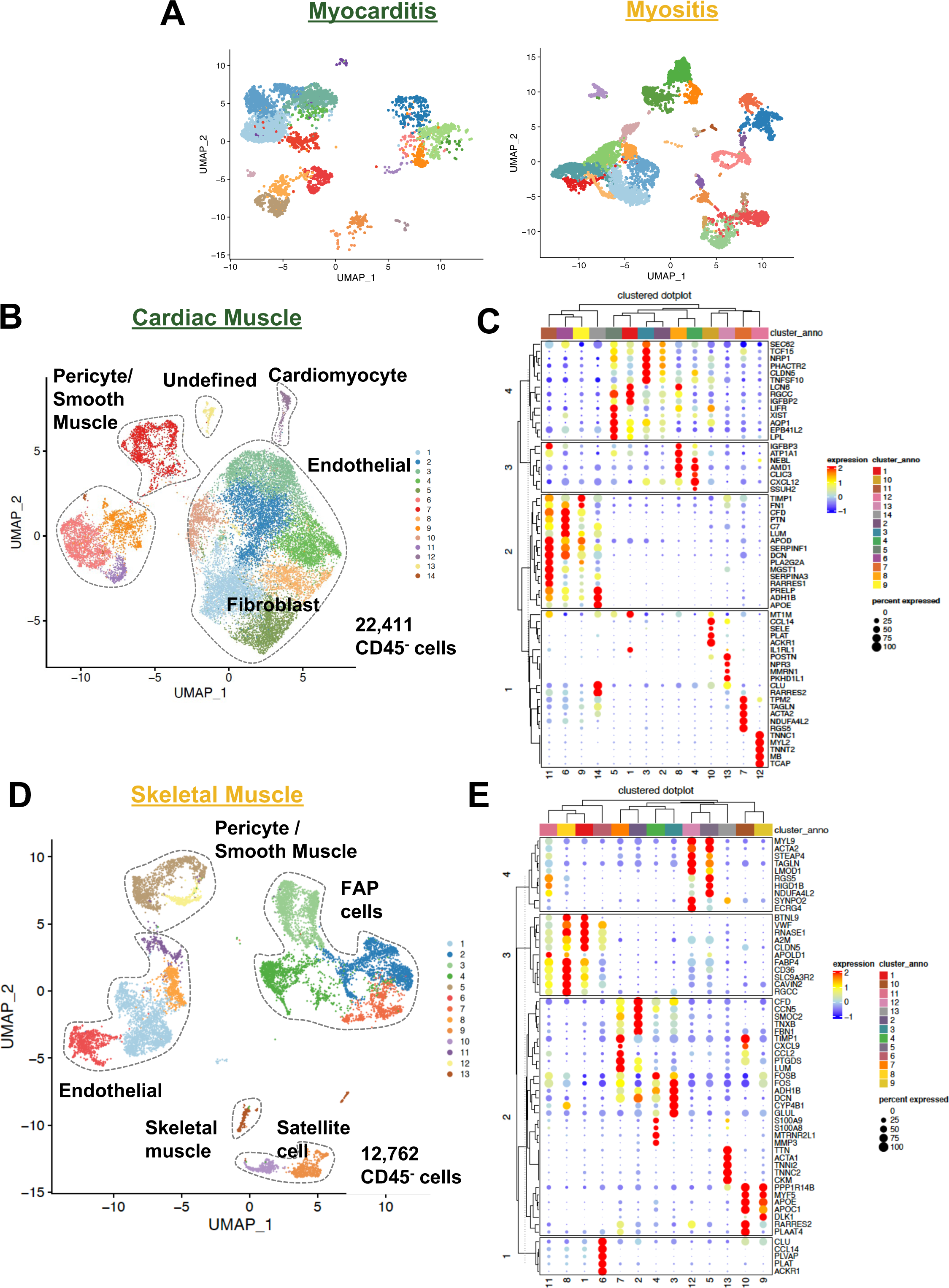
Landscape of CD45^-^ (non-immune) populations from patient cardiac and skeletal muscle tissues. **A.** UMAP projections of myocarditis (left) and myositis (samples) downsampled in toxicity groups to match cell counts with control samples. **B.** UMAP projections of CD45^-^ cells in cardiac muscle. **C.** Dot plots indicating cluster definitions for CD45^-^ cells in cardiac muscle. **D.** UMAP projections of CD45^-^ cells in skeletal muscle. **E.** Dot plots indicating cluster definitions for CD45^-^ cells in skeletal muscle.

**Supplementary Figure 3.**
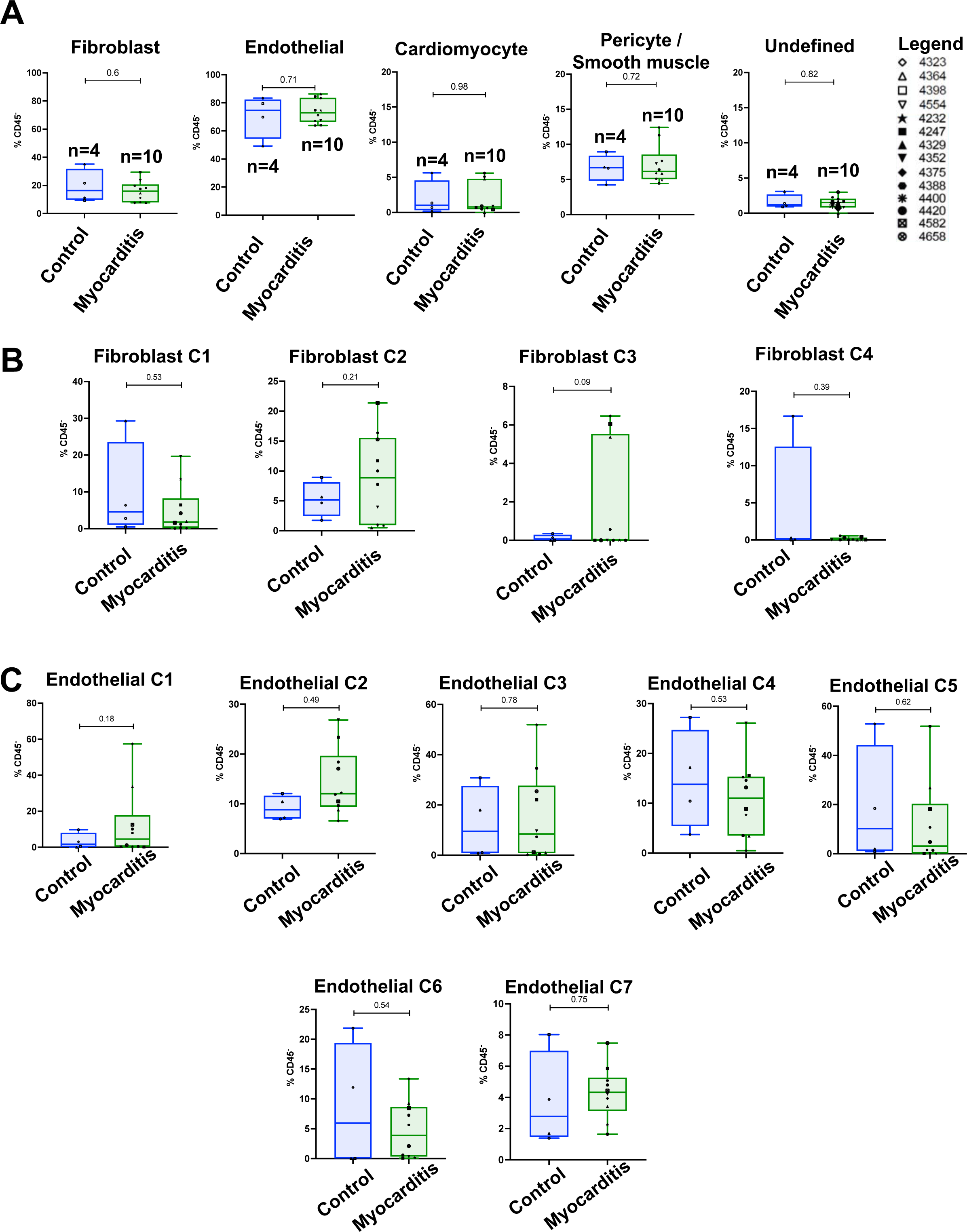
Analysis of CD45^-^ subpopulations in cardiac muscle. **A.** Frequency of CD45^-^ cell clusters in immune-related myocarditis versus control cardiac muscle tissues. **B.** Frequency of fibroblast subsets in cardiac muscle. **C.** Frequency of endothelial cell subsets in cardiac muscle.

**Supplementary Figure 4.**
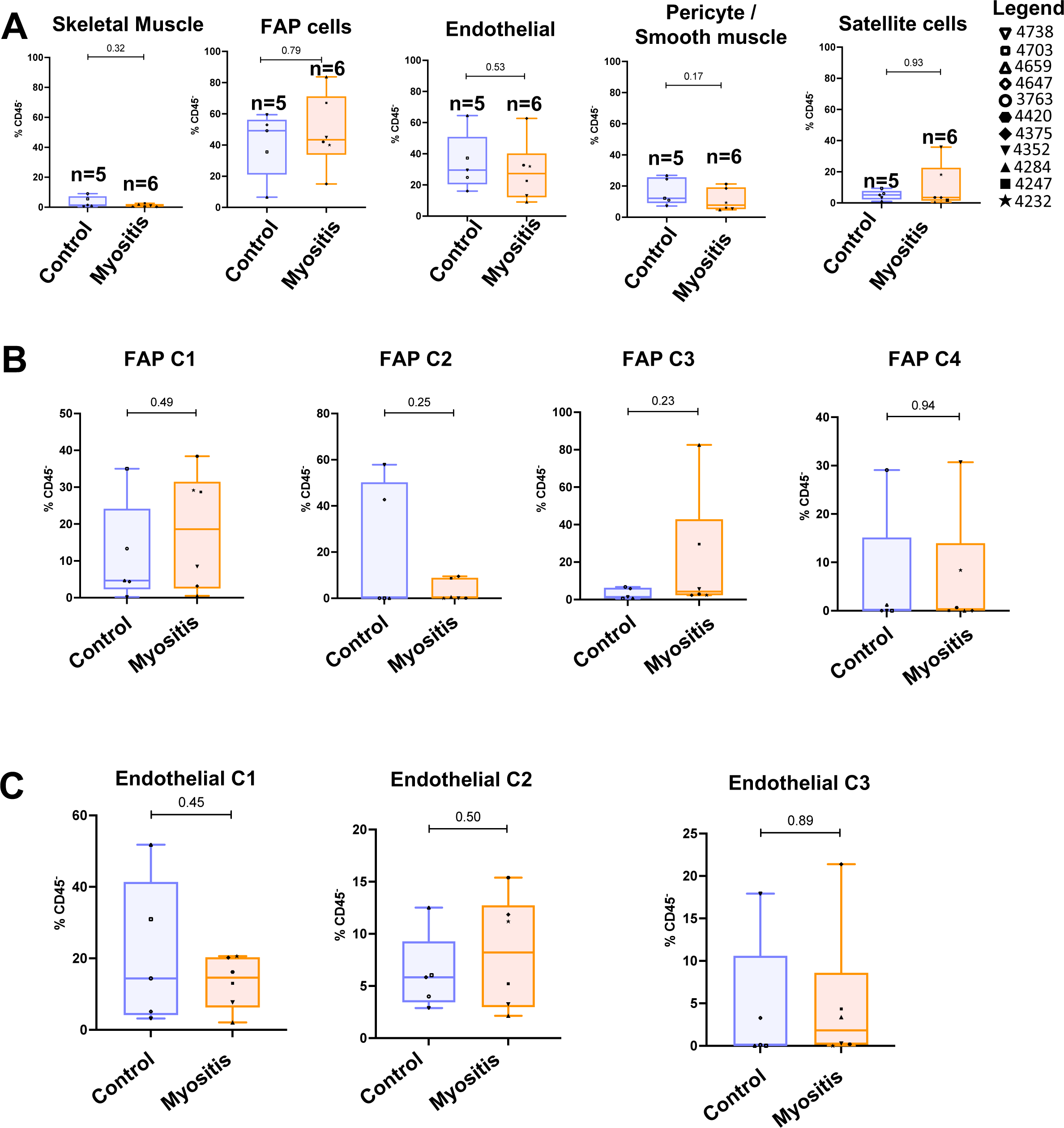
Analysis of CD45^-^ subpopulations in cardiac muscle. **A.** Frequency of CD45-cell clusters in immune-related myositis versus control skeletal muscle tissues. **B.** Frequency of fibroadipogenic precursor (FAP) cell subsets in cardiac muscle. **C.** Frequency of endothelial cell subsets in cardiac muscle.

**Supplementary Figure 5.**
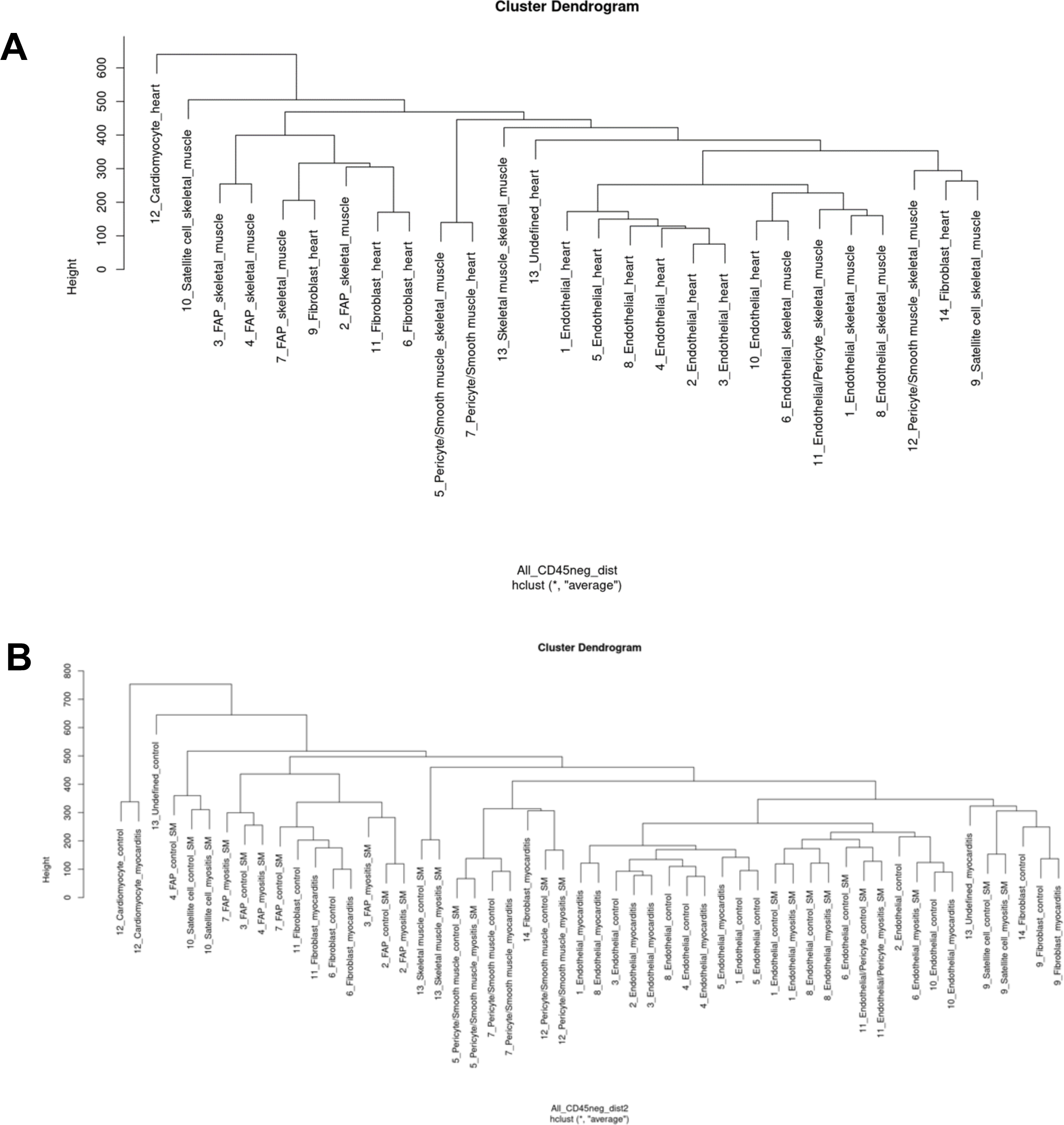
Hierarchical clustering based on Euclidean distance of expression in CD45^-^ cells. **A.** Cluster dendrogram of all cardiac muscle and all skeletal muscle samples. **B.** Cluster dendrogram of myocarditis, control cardiac muscle, myositis, and control skeletal muscle samples.

**Supplementary Figure 6:**
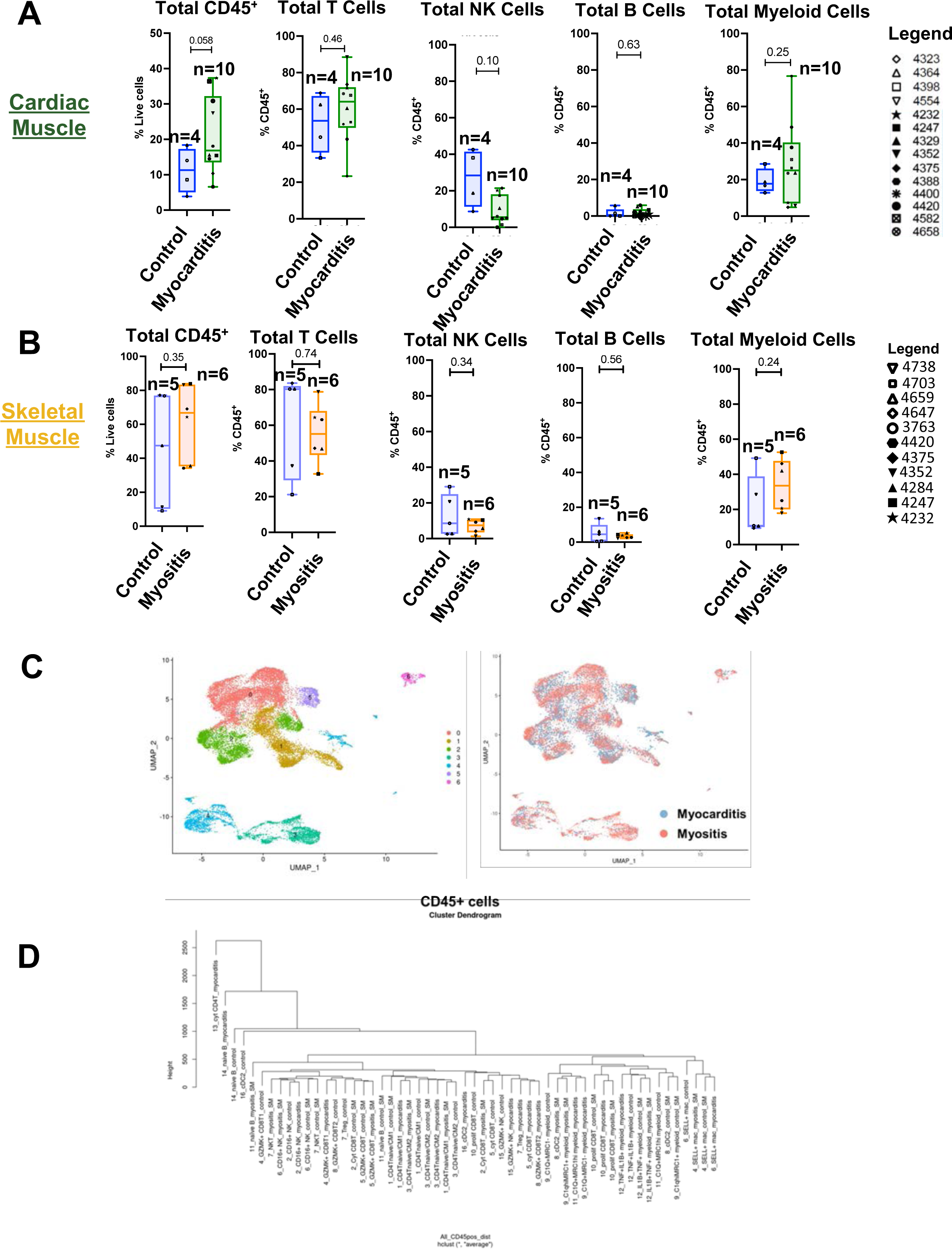
Analysis of total CD45^+^ immune cell populations. **A.** Frequency of CD45^+^ cells and total immune cell subsets in cardiac muscle from patients with myocarditis compared with controls. **B.** Frequency of CD45^+^ cells and total immune cell subsets in skeletal muscle from patients with myositis compared with controls. **C.** UMAP overlay of CD45^+^ cells in myocarditis and myositis samples. **D.** Cluster dendrogram of CD45+ cells in myocarditis, control cardiac muscle, myositis, and control skeletal muscle samples.

**Supplementary Figure 7.**
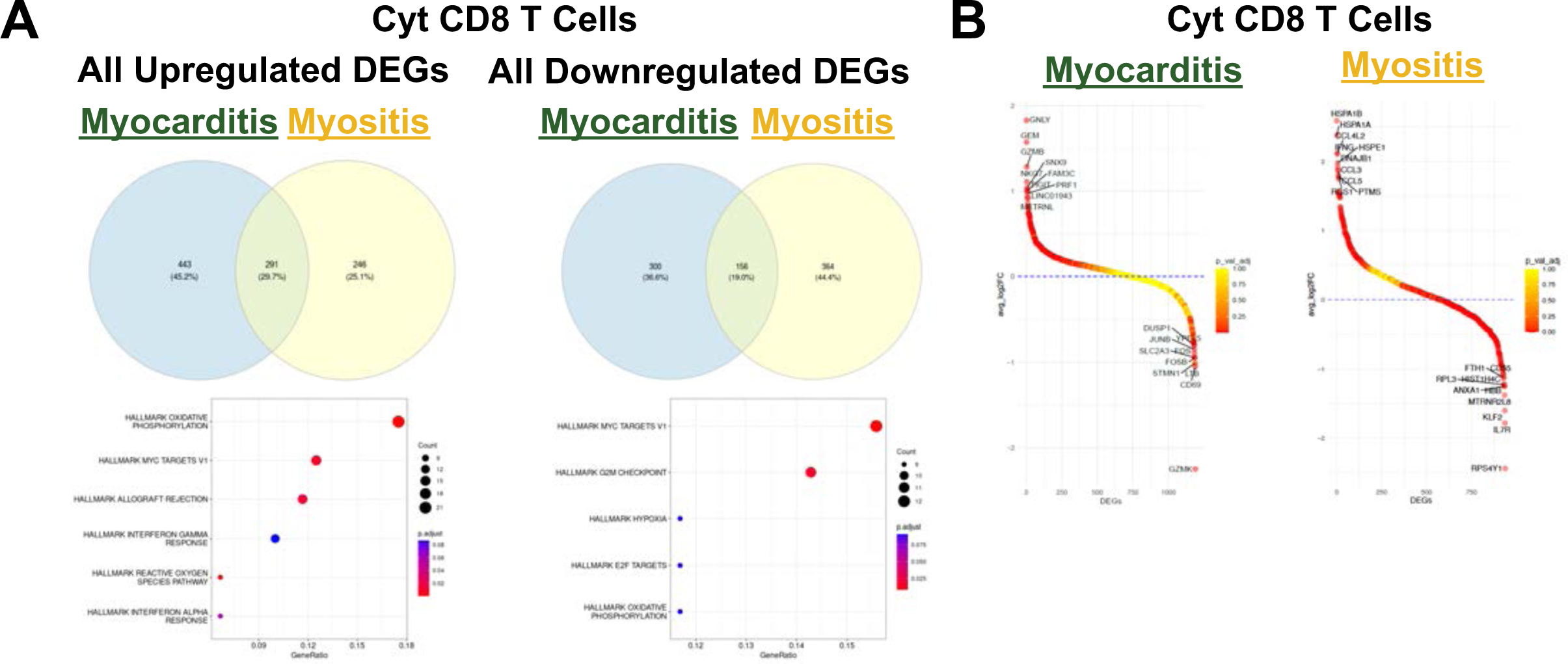
Differentially expressed genes (DEGs) in Cytotoxic CD8^+^ T Cells. **A.** Comparison of percentages of upregulated and downregulated DEGs shared and distinct within cytotoxic CD8^+^ T cell populations in myocarditis versus myositis samples. **B.** Identity of most highly upregulated and downregulated genes within cytotoxic CD8^+^ T cell populations in myocarditis and myositis samples.

**Supplementary Figure 8.**
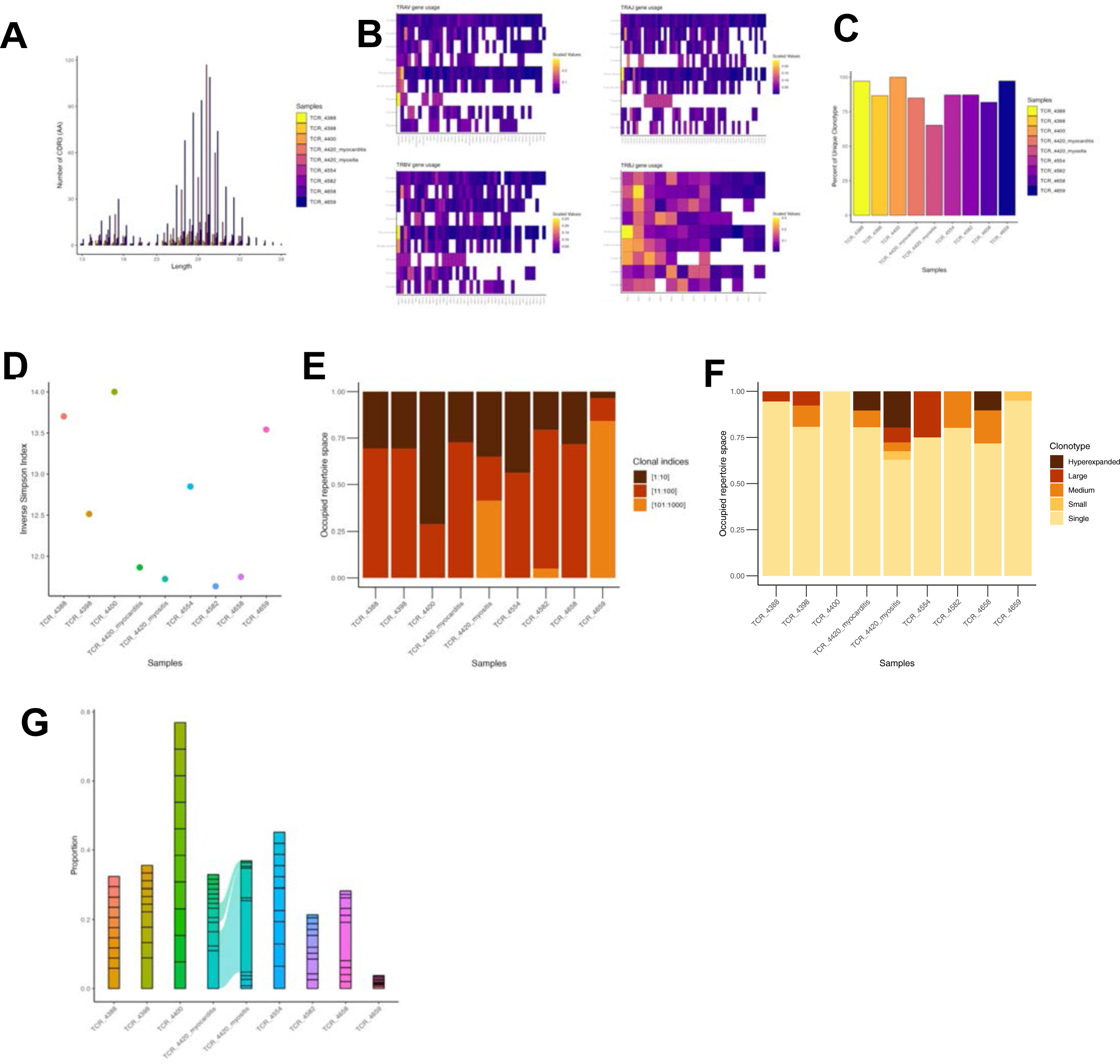
Single cell TCR sequencing (scRNASeq) from patient cardiac and skeletal muscle tissue. **A.** Distribution of CDR3 length. **B.** V and J gene usage. **C.** Clonotypes defined based on V(D)J gene usage and CDR3 nucleotide sequence. **D.** TCR clonal diversity assessed by inverse Simpson methods. **E.** Occupied receptor scores. **F.** TCR clonal expansion. **G**. Alluvial plot showing the overlapping of top 10 TCR clones in each sample.

**Supplementary Figure 9.**
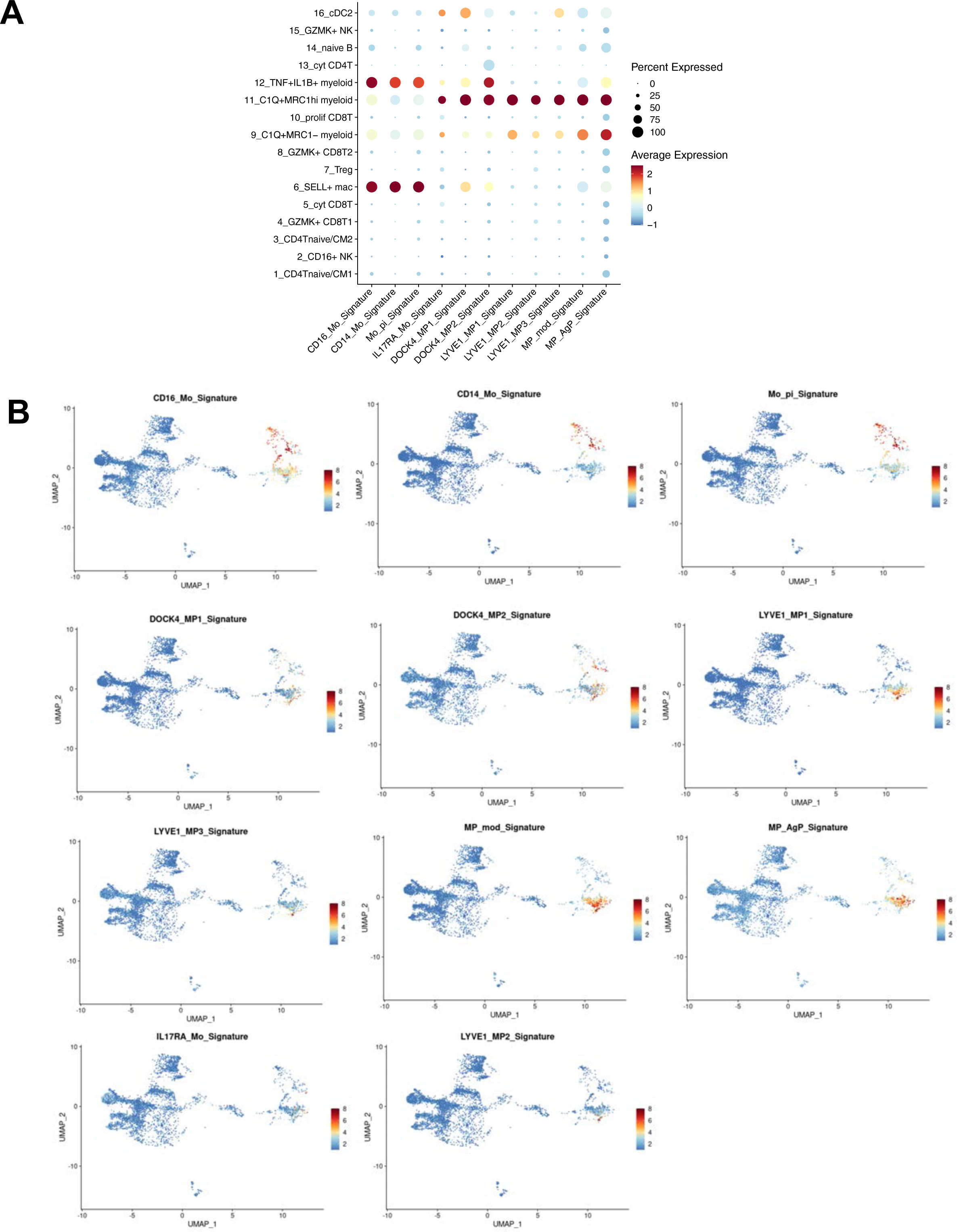
Comparison of myeloid cell subclusters in cardiac muscle with previously reported subsets in human heart. **A.** Comparison of myeloid subclusters with signature score derived from top 30 differentially expressed genes from previously reported myeloid cell subsets. **B.** Feature plots indicating presence of previously reported myeloid cell subsets in cardiac muscle samples.

**Supplementary Figure 10.**
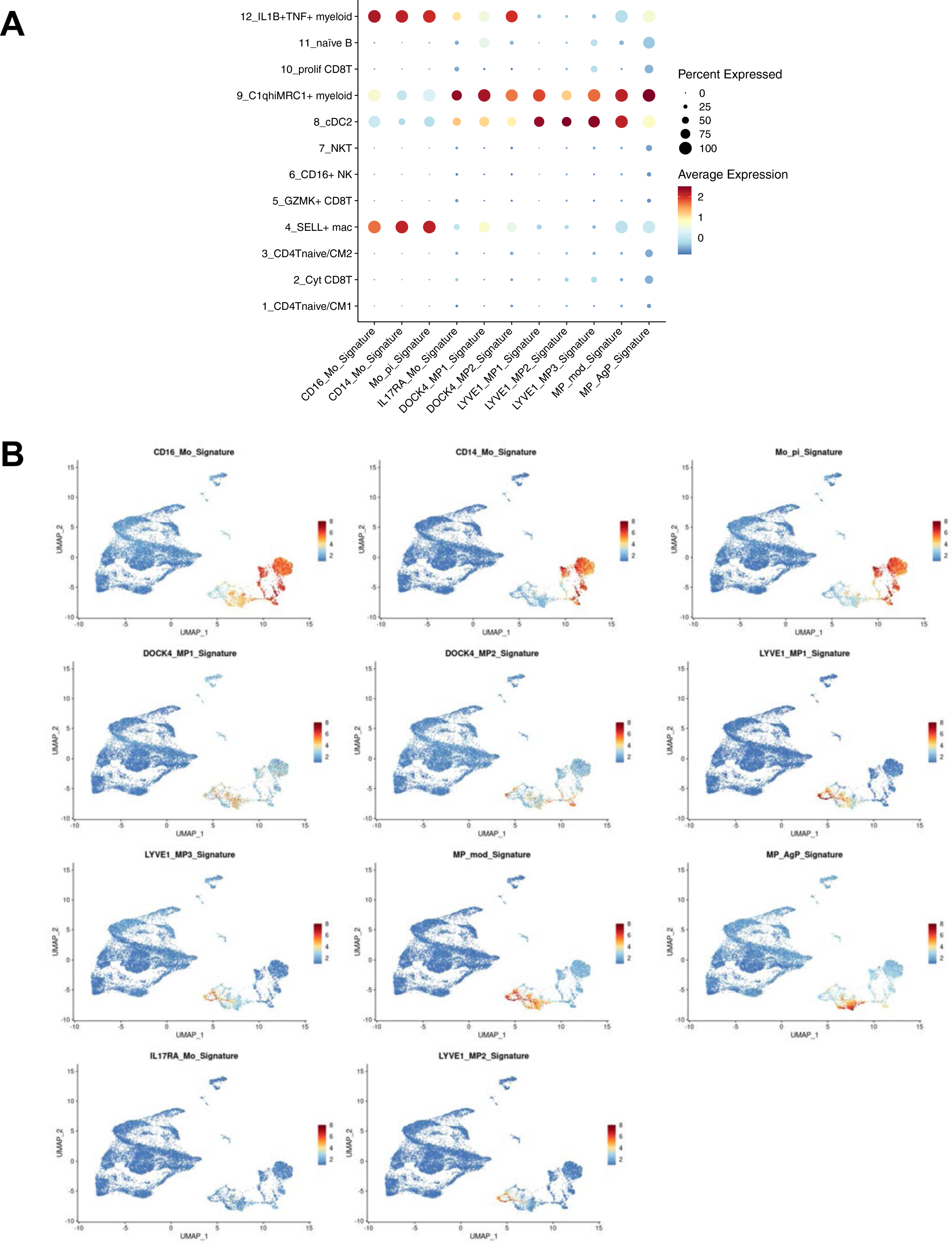
Comparison of myeloid cell subclusters in skeletal muscle with previously reported subsets. **A.** Comparison of myeloid subclusters with signature score derived from top 30 differentially expressed genes from previously reported myeloid cell subsets. **B.** Feature plots indicating presence of previously reported myeloid cell subsets in skeletal muscle samples.

**Supplementary Figure 11.**
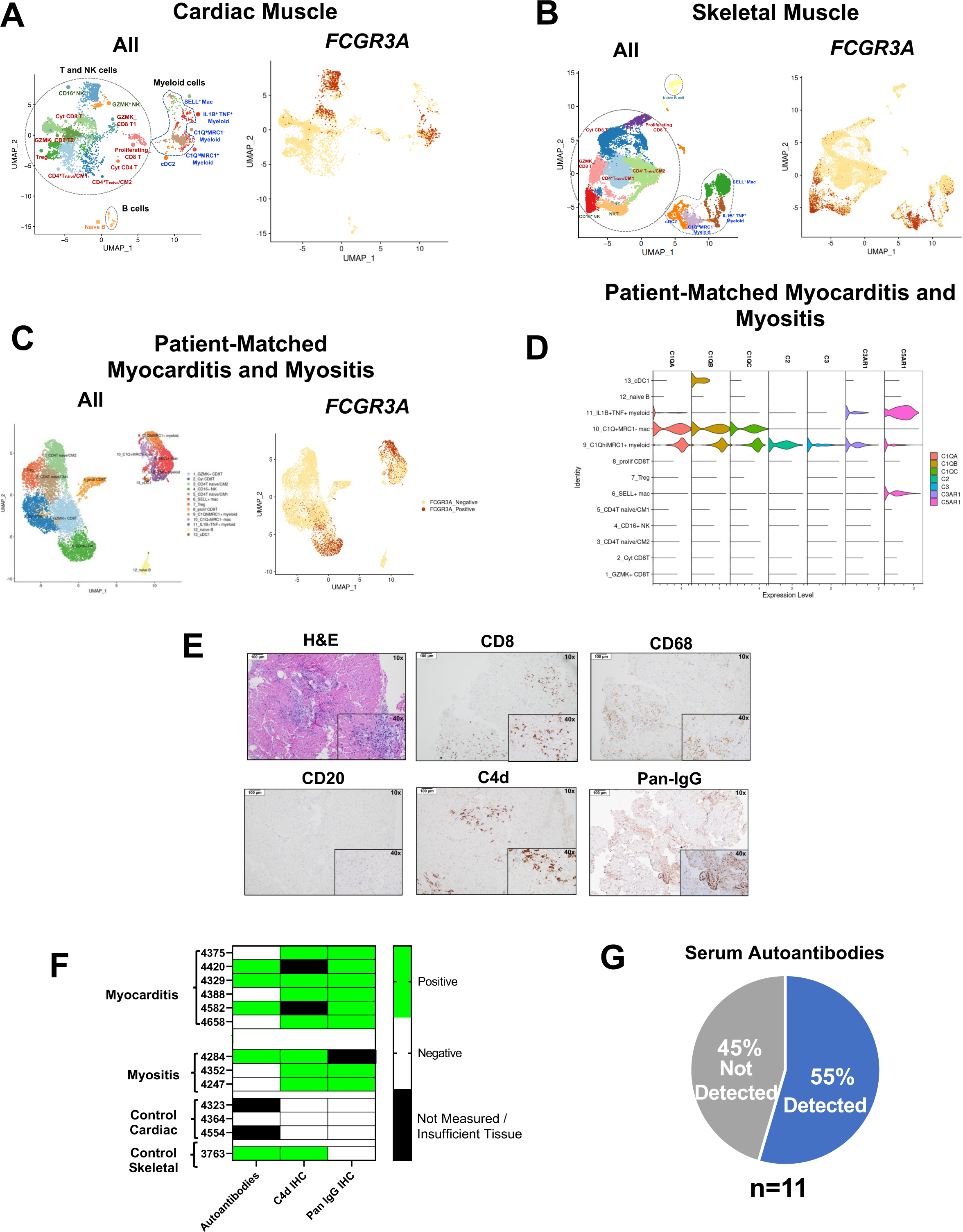
Complement activation associated with serum autoantibodies and tissue IgG deposition in ICT-induced myocarditis and myositis. **A.** Feature plot of *FCGR3A*-expressing cells in cardiac muscle, with all subclusters shown for reference (left). **B.** Feature plot of *FCGR3A*-expressing cells in skeletal muscle, with all subclusters shown for reference (left). **C.** Feature plot of *FCGR3A*-expressing cells in patient-matched myocarditis and myositis samples, with all subclusters shown for reference (left). **D.** Violin plot of expression of classical complement pathway associated genes in immune cell subsets in patient-matched cardiac and skeletal muscle samples. **E.** Representative hematoxylin and eosin (H&E) and immunohistochemistry (IHC) for T cells (CD4, CD8), myeloid cells (CD68), B cells (CD20), pan-IgG and C4d in cardiac muscle from patient with immune-related myocarditis. All images shown at 10x magnification, with inset at 40x magnification. All stains are taken from patient #4329. **F**. Heat map evaluating correlation of serum autoantibodies and IHC for C4d and Pan-IgG in cardiac and skeletal muscle samples. **G.** Distribution of detectable striated muscle or acetylcholine receptor binding or modulating IgGs in serum.

**Supplementary Figure 12.**
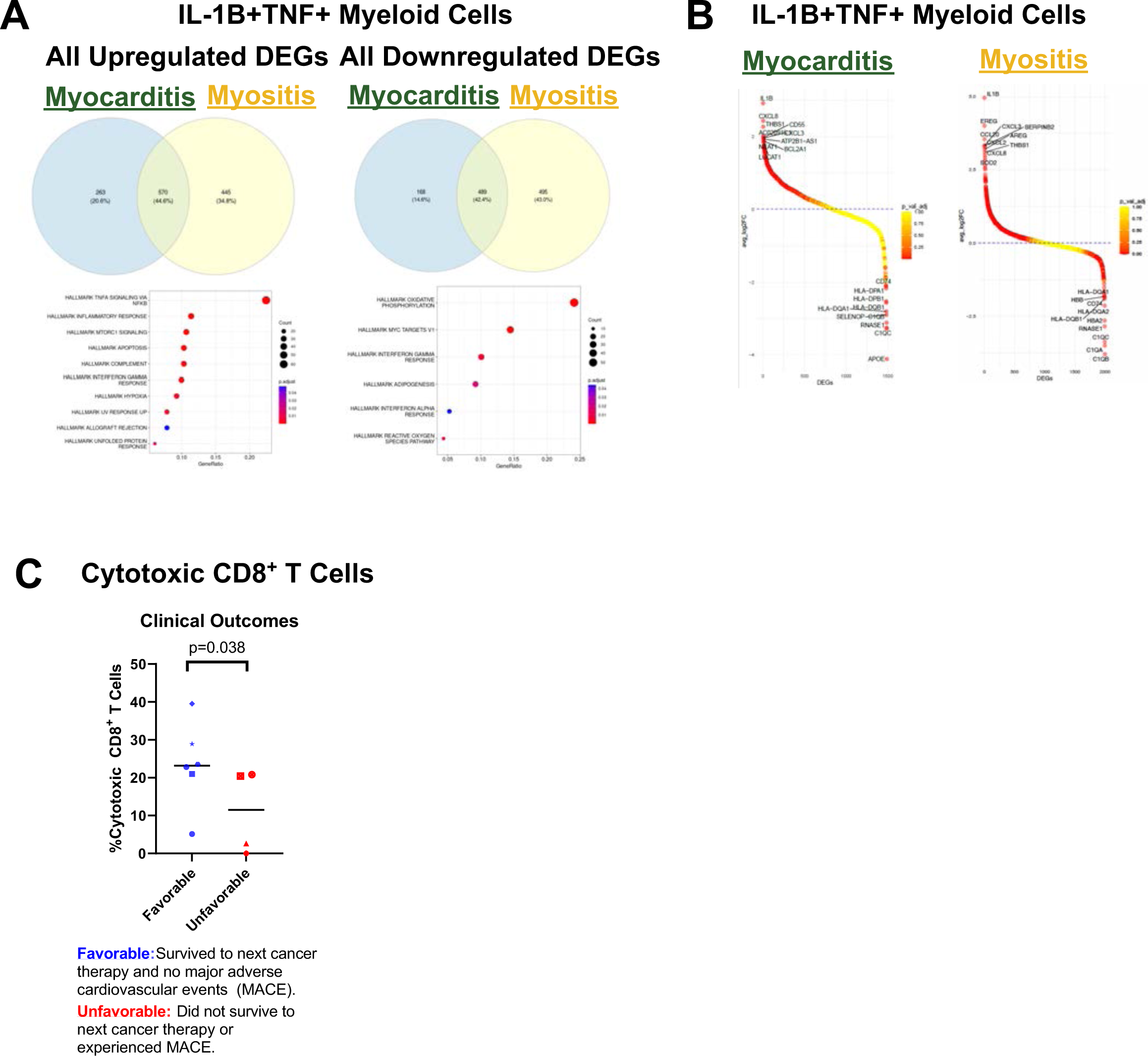
Differentially expressed genes (DEGs) in IL1B^+^ Myeloid cell subsets and association of frequency of cytotoxic CD8^+^ T cells with clinical outcomes. **A.** Comparison of percentages of upregulated and downregulated DEGs shared and distinct within IL-1B^+^TNF^+^ myeloid populations in myocarditis versus myositis samples. **B.** Identity of most highly upregulated and downregulated genes within IL-1B^+^TNF^+^ myeloid populations in myocarditis and myositis samples. **C.** Association of frequency of cytotoxic CD8^+^ T cells with clinical outcomes.

**Supplemental Item 1:**
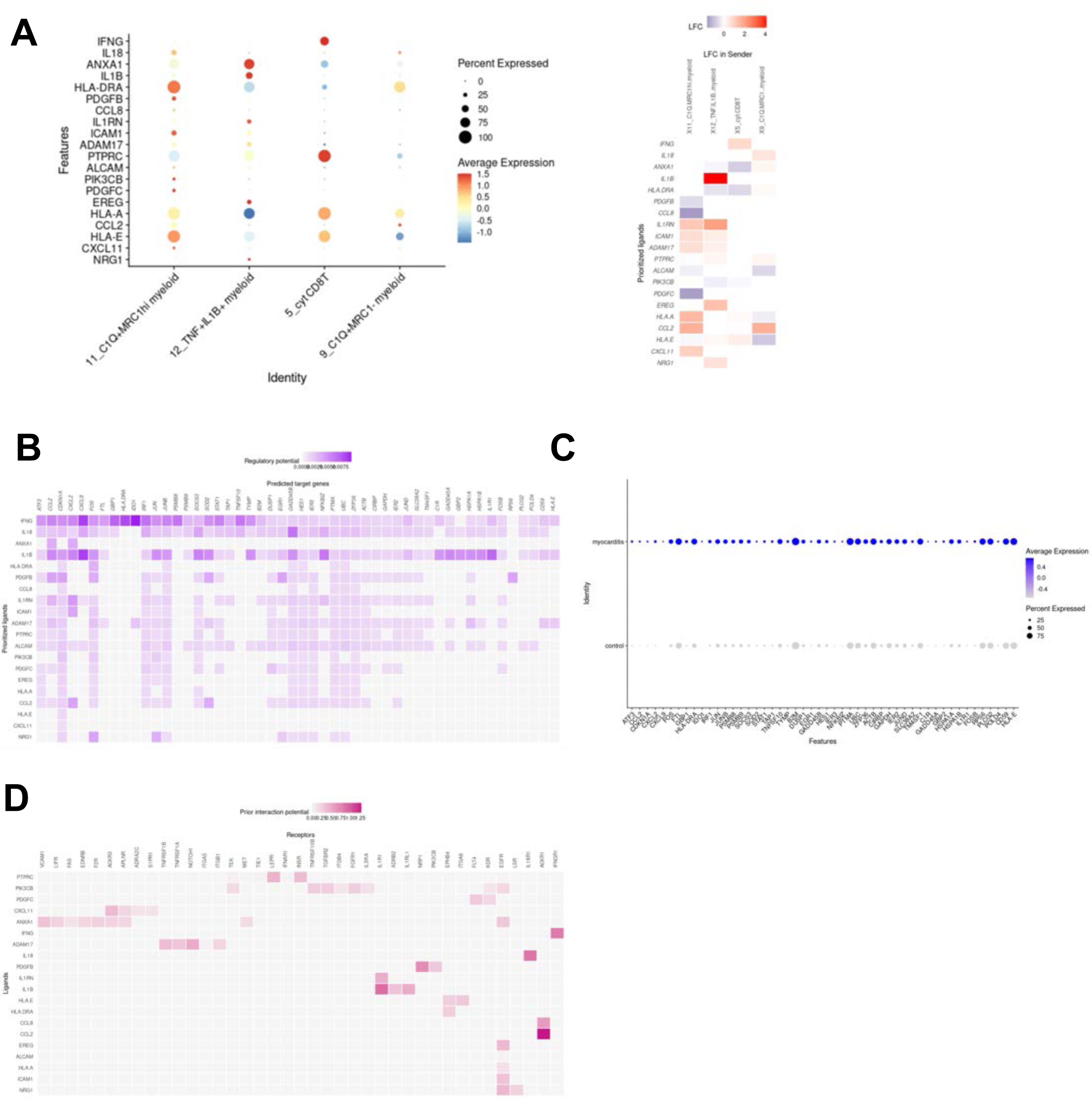
Cell-cell communication analysis from myeloid cells and cytotoxic T cells to endothelial cells in myocarditis. **A.** NicheNet analysis analyzing communication from myeloid cells and cytotoxic CD8^+^ T cells to endothelial cells in myocarditis versus control samples. Left: Expression of active ligands in sender cells; Right: Prioritized ligands by Pearson correlation coefficient. **B.** Expression of predicted target genes from the prioritized ligands from myeloid cells and cytotoxic T cells (sender cells) to endothelial cells (receiver cells). **C**. Increased expression of complement gene *C1R* and stress-related response genes in endothelial cells in myocarditis versus control samples **D.** Inferred ligand-receptor interactions for top-ranked ligands.

**Supplemental Item 2:**
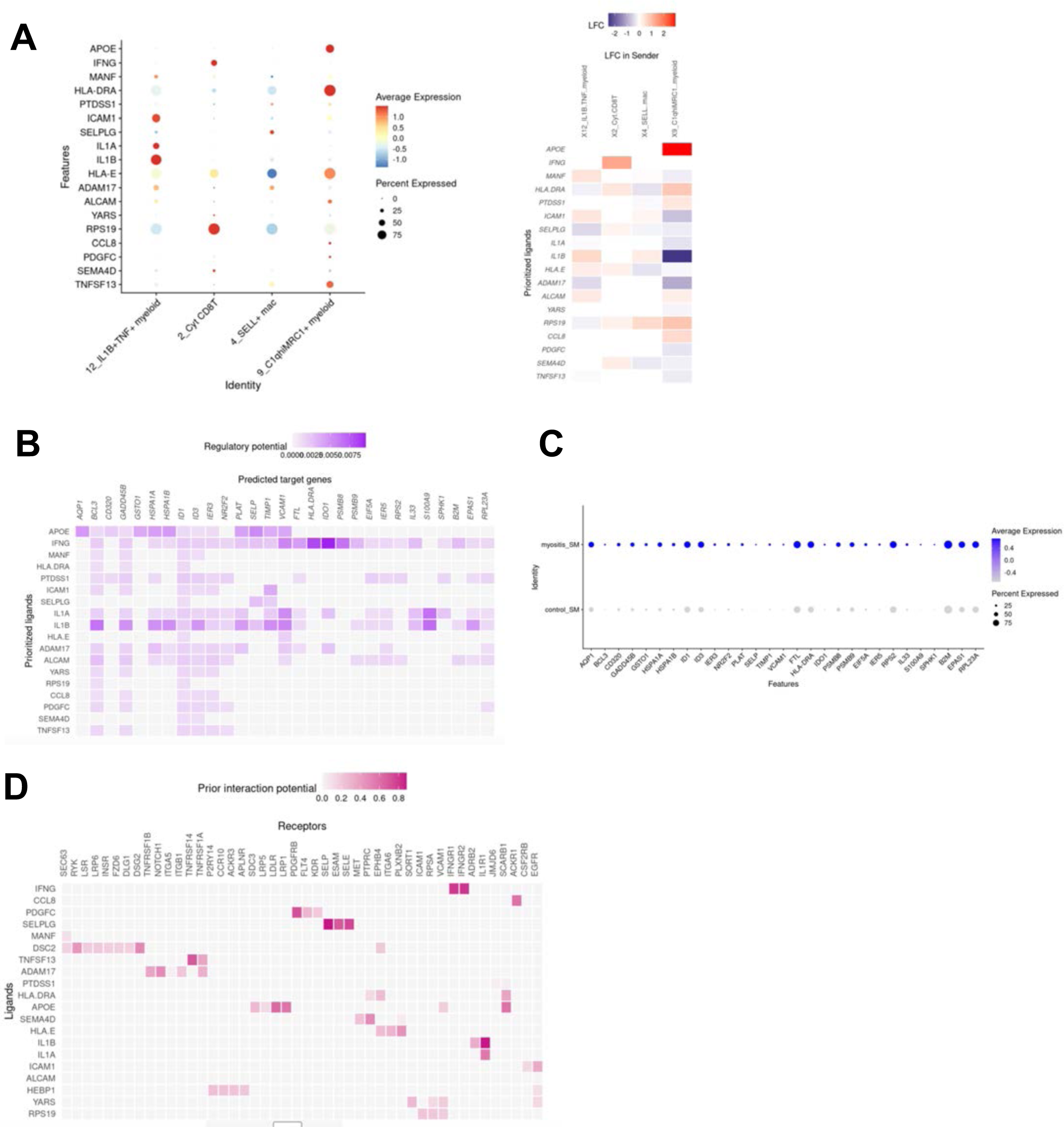
Cell-cell communication analysis from myeloid cells and cytotoxic T cells to endothelial cells in myositis. **A.** NicheNet analysis analyzing communication from myeloid cells and cytotoxic CD8^+^ T cells to endothelial cells in myositis versus control samples. Left: Expression of active ligands in sender cells; Right: Prioritized ligands by Pearson correlation coefficient. **B.** Expression of predicted target genes from the prioritized ligands from myeloid cells and cytotoxic T cells (sender cells) to endothelial cells (receiver cells). **C**. Increased expression of stress-related response genes in endothelial cells in myositis versus control samples **D.** Inferred ligand-receptor interactions for top-ranked ligands.

**Supplemental Item 3:**
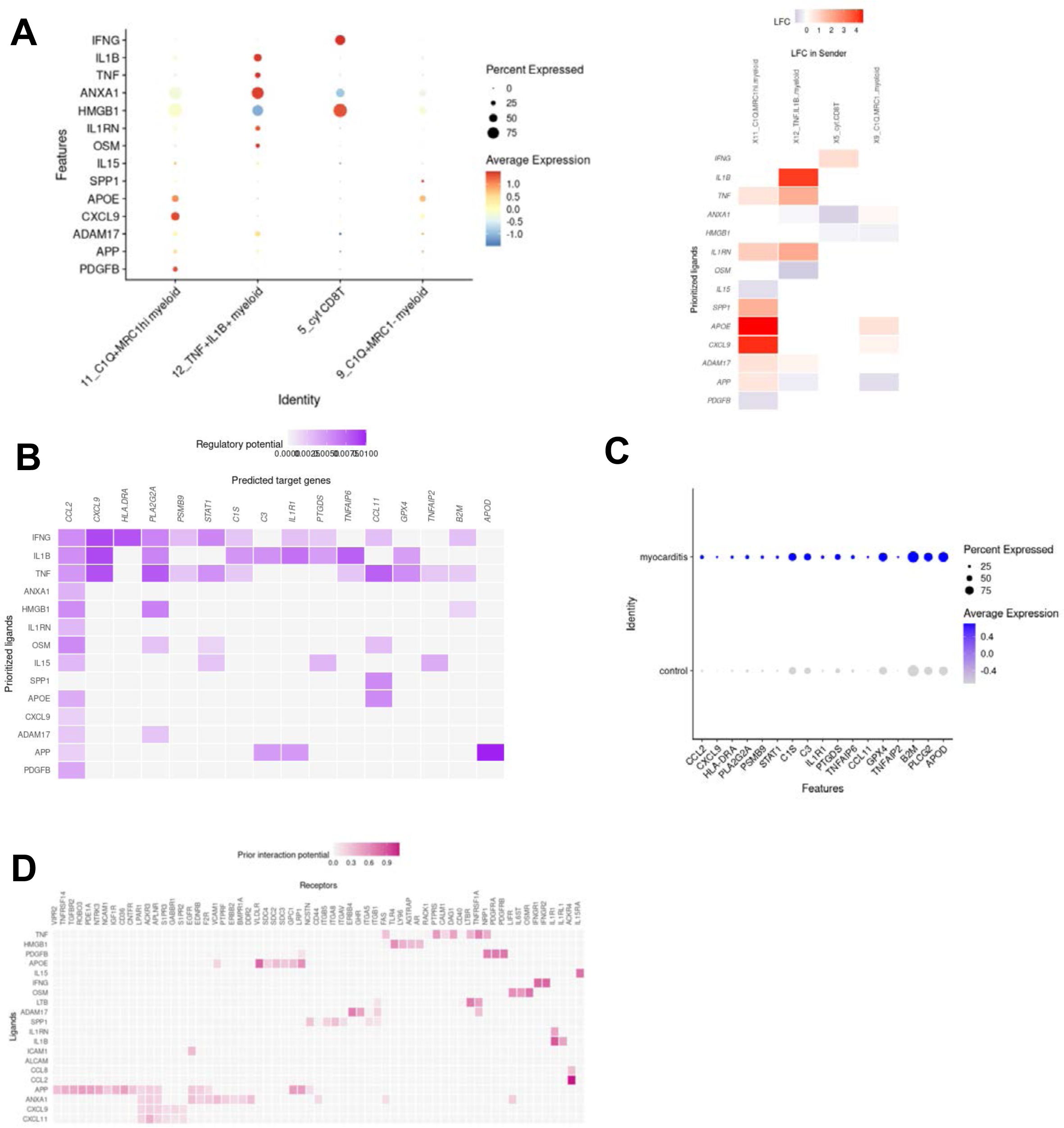
Cell-cell communication analysis from myeloid cells and cytotoxic T cells to fibroblasts in myocarditis. **A.** NicheNet analysis analyzing communication from myeloid cells and cytotoxic CD8^+^ T cells to fibroblasts in myocarditis versus control samples. Left: Expression of active ligands in sender cells; Right: Prioritized ligands by Pearson correlation coefficient. **B.** Expression of predicted target genes from the prioritized ligands from myeloid cells and cytotoxic T cells (sender cells) to fibroblasts (receiver cells). **C**. Increased expression of complement genes *C3* and *C1S* in myocarditis versus control samples downstream of IL-1B signaling. **D.** Inferred ligand-receptor interactions for top-ranked ligands. **E.** Representative hematoxylin and eosin (H&E) and immunohistochemistry (IHC) for CD8 T cells, CD20 B cells, CD68 myeloid cells, C4d, and pan-IgG in cardiac muscle from patient #4329 with immune-related myocarditis. All images shown at 10x magnification, with inset at 40x magnification. **F.** Semiquantitative correlation of C4d and Pan IgG IHC in cardiac tissue with presence of autoantibodies (0: no staining; 1+ moderate staining; 2+ intense staining).

**Supplemental Item 4.**
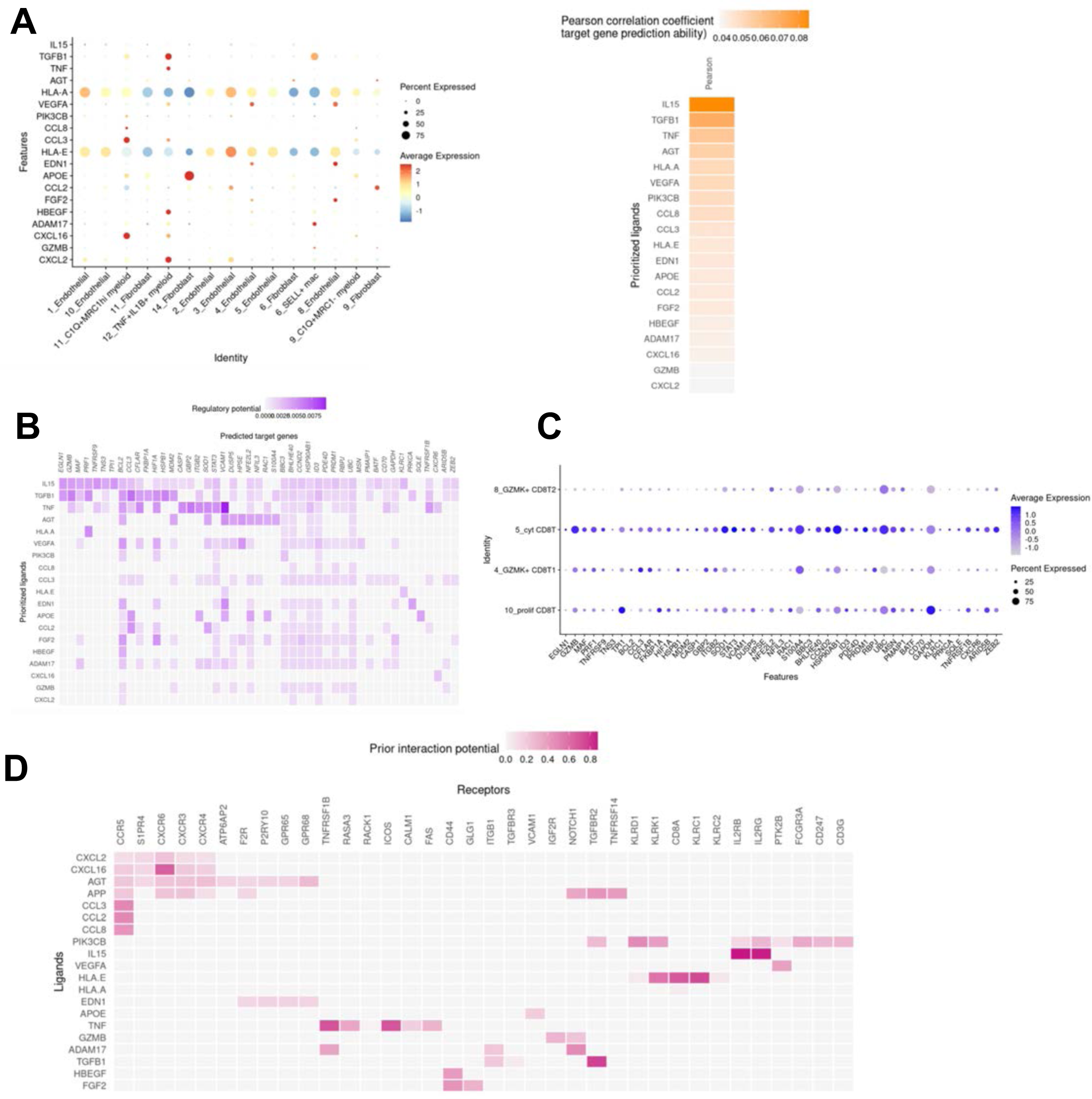
Cell-cell communication analysis from fibroblasts, endothelial cells, and myeloid cells to cytotoxic T cells in myocarditis. **A.** NicheNet analysis analyzing communication from fibroblasts, endothelial cells, myeloid cells to cytotoxic CD8 T cells compared to other CD8 T cell clusters myocarditis. Left: Expression of active ligands in sender cells; Right: Prioritized ligands by Pearson correlation coefficient. **B.** Expression of predicted target genes from the prioritized ligands from fibroblasts, endothelial cells, and myeloid cells (sender cells) to cytotoxic T cells cells (receiver cells). **C**. Relative gene expression in cytotoxic CD8 T cells compared to other TCD8 T cell clusters. **D.** Inferred ligand-receptor interactions for top-ranked ligands.

**Supplemental Item 5.**
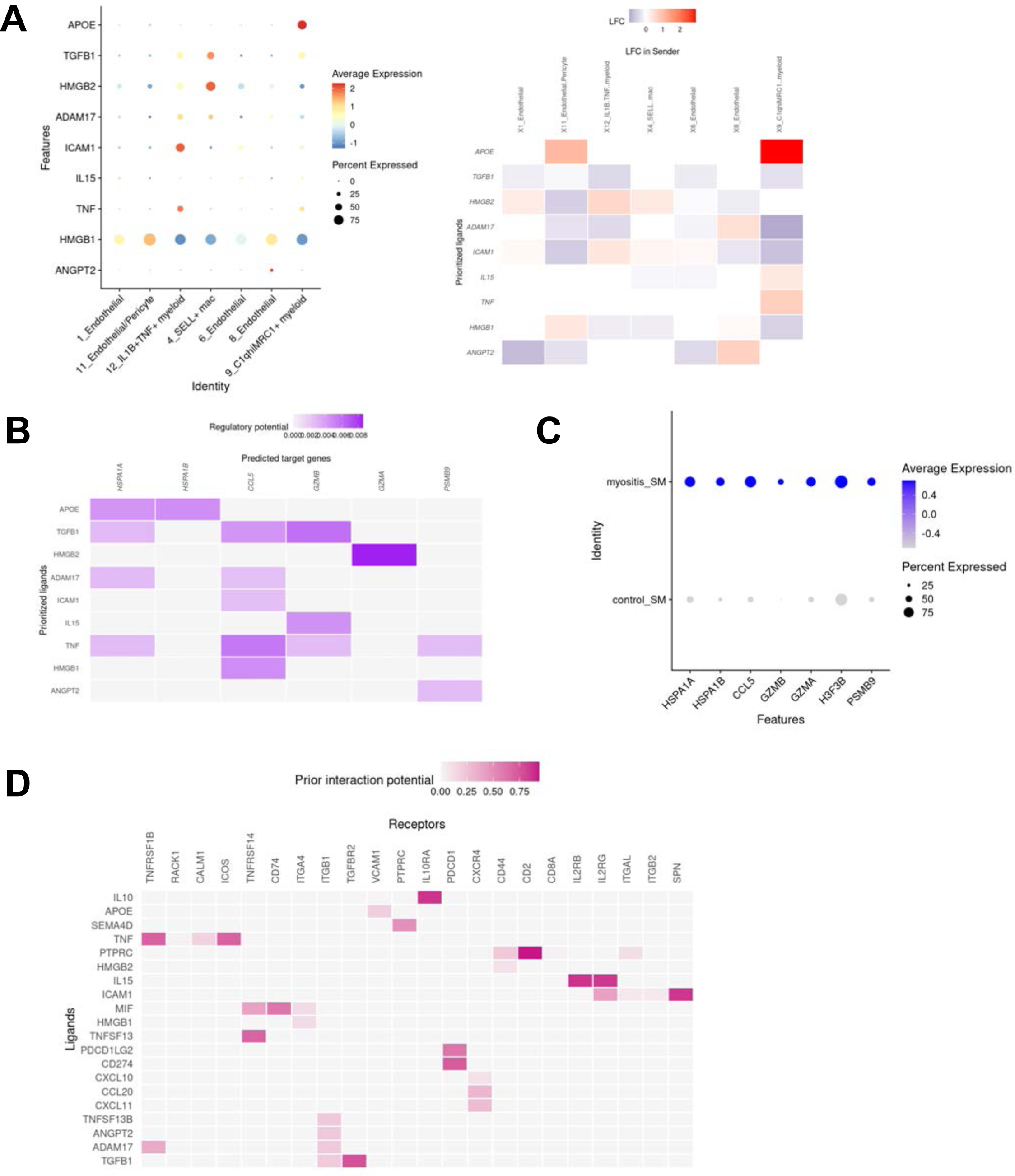
Cell-cell communication analysis from endothelial cells and myeloid cells to cytotoxic T cells in myositis. **A.** NicheNet analysis analyzing communication from endothelial cells and myeloid cells to cytotoxic CD8 T cells in myositis. Left: Expression of active ligands in sender cells; Right: Prioritized ligands by Pearson correlation coefficient. **B.** Expression of predicted target genes from the prioritized ligands from endothelial cells and myeloid cells (sender cells) to cytotoxic T cells cells (receiver cells). **C**. Relative gene expression in cytotoxic CD8 T cells compared to other TCD8 T cell clusters. **D.** Inferred ligand-receptor interactions for top-ranked ligands.

**Supplementary Table 1:**
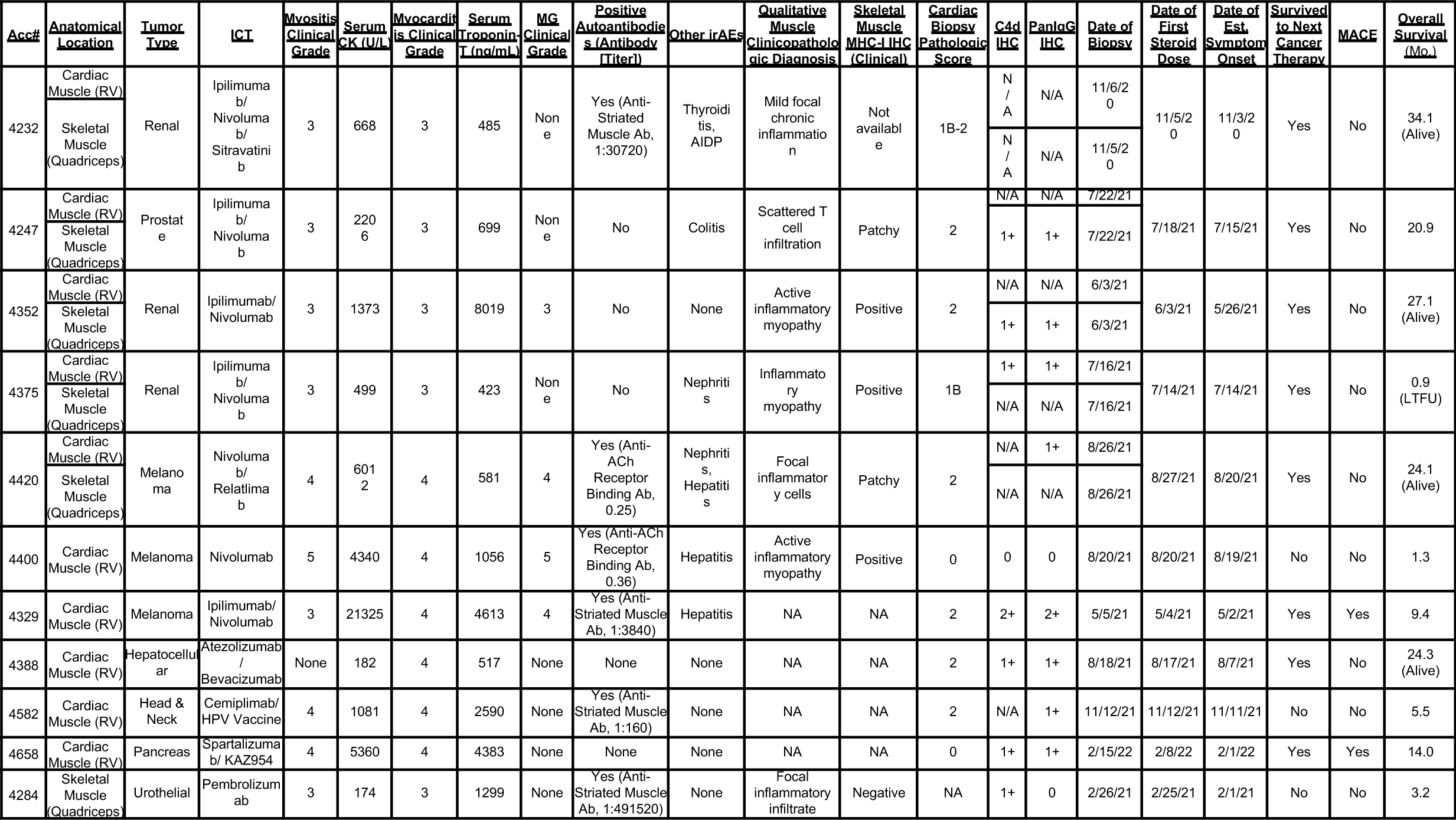
Clinical characteristics of individual patients with immune-related myocarditis and/or myositis. AIDP=acute inflammatory demyelinating polyneuropathy. CK=creatine kinase; MG=myasthenia gravis; RV=right ventricle. Reference ranges: CK <180 U/L; troponin-T <18 ng/mL. MACE=major adverse cardiovascular events. LTFU=lost to follow-up

**Supplementary Table 2:**
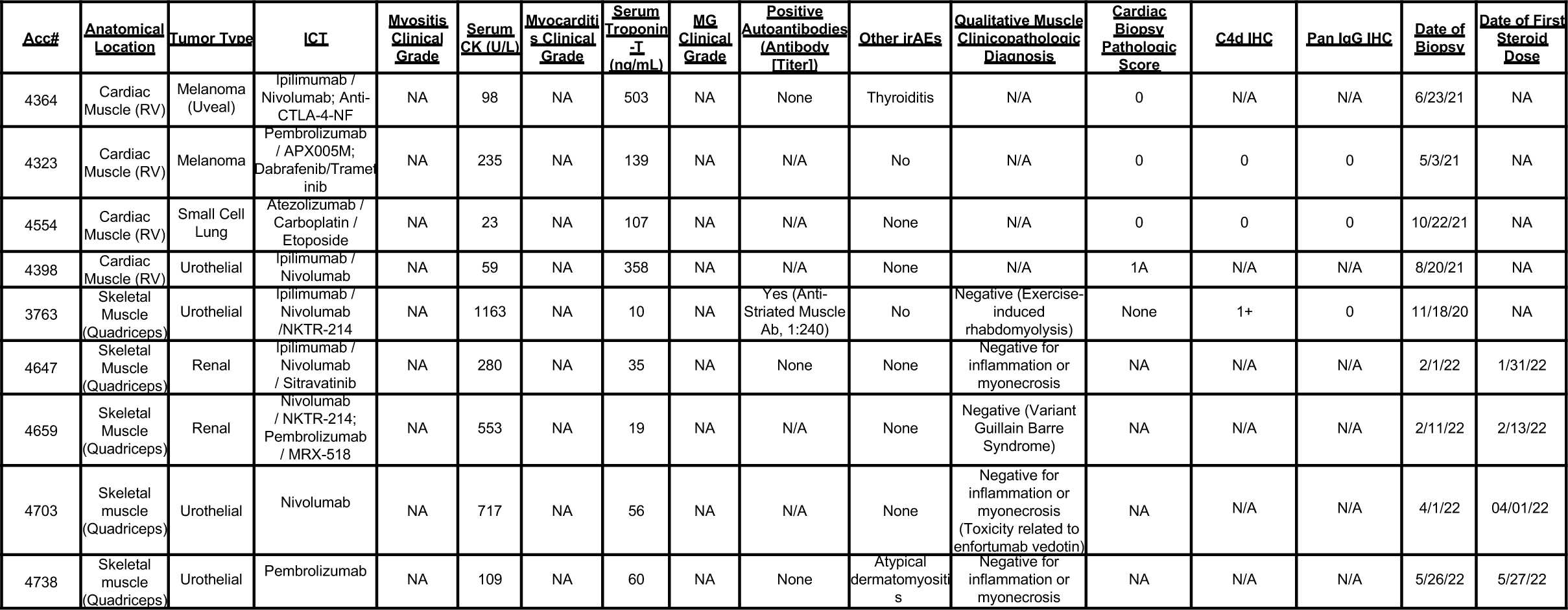
Clinical characteristics of control patients. CK=creatine kinase; MG=myasthenia gravis; RV=right ventricle. Reference ranges: CK <180 U/L; troponin-T <18 ng/mL.

